# Oscillations in spontaneous and visually evoked neuronal activity in the superficial layers of the cat’s superior colliculus: Oscillations in neuronal activity in superficial SC

**DOI:** 10.1101/347799

**Authors:** Andrzej T. Foik, Anaida Ghazaryan, Wioletta J. Waleszczyk

## Abstract

Oscillations are ubiquitous features of neuronal activity in sensory systems and are considered as a substrate for the integration of sensory information. Several studies have described oscillatory activity in the geniculate visual pathway, but little is known about this phenomenon in the extrageniculate visual pathway. We describe oscillations in evoked and background activity in the cat’s superficial layers of the superior colliculus, a retinorecipient structure in the extrageniculate visual pathway. Extracellular singleunit activity was recorded during periods with and without visual stimulation under isoflurane anesthesia in the mixture of N_2_O/O_2_. Autocorrelation, FFT and renewal density analyses were used to detect and characterize oscillations in the neuronal activity. Oscillations were common in the background and stimulus-evoked activity. Frequency range of background oscillations spanned between 5 – 90 Hz. Oscillations in evoked activity were observed in about half of the cells and could appear in two forms – stimulus-phase-locked (10 – 100 Hz), and stimulus-phase-independent (8 – 100 Hz) oscillations. Stimulus-phase-independent and background oscillatory frequencies were very similar within activity of particular neurons suggesting that stimulus-phase-independent oscillations may be a form of enhanced “spontaneous” oscillations. Stimulus-phase-locked oscillations were present in responses to moving and flashing stimuli. In contrast to stimulus-phase-independent oscillations, the strength of stimulus-phase-locked oscillations was positively correlated with stimulus velocity and neuronal firing rate. Our results suggest that in the superficial layers of the superior colliculus stimulus-phase-independent oscillations may be generated by the same mechanism(s) that lie in the base of “spontaneous” oscillations, while stimulus-phase-locked oscillations may result from interactions within the intra-collicular network and/or from a phase reset of oscillations present in the background activity.

## Introduction

Information about the external world is transferred to the brain via trains of action potentials that can exhibit a repetitive temporal structure. These temporal patterns, formed by the periodic appearance of single spikes, bursts of spikes, or periodic changes in the firing rate are called oscillations (Lestienne, 2001). Oscillations of certain neuronal groups are characteristic of different behavioral states (for revs. see Buzaki, 2006; Singer, 2009; Uhlhaas et al., 2009). Many studies have described oscillatory activity in the geniculo-cortical visual pathway in different species and substantial knowledge has been accumulated about the characteristics, origin, stimulus dependency, and the role of oscillatory neuronal activity in the structures of this pathway (Eckhorn et al. 1988; Gray et al. 1989, 1990; Eckhorn and Obermueller, 1993; Molotchnikoff et al. 1996; Bringuier et al. 1997; Castelo-Branco et al., 1998; Kohn and Smith, 2005; Samonds and Bonds, 2005; Fries, 2009; Koepsell, 2009; 2010; Ito et al., 2010; Lima et al., 2010), but little is known about this phenomenon in the extrageniculate visual pathway.

The extrageniculate pathway transmits visual information to higher (association) visual cortices via optic tectum and other thalamus areas thus effectively bypassing the lateral geniculate nucleus (LGN). The superior colliculus (SC), a major retinorecipient structure of the midbrain, is the first structure in this pathway, which transfers visual information to the cortex via thalamic nuclei (lateral posterior-pulvinar complex and suprageniculate nucleus of the posterior thalamus (Graybiel and Berson, 1980; Abramson and Chalupa 1988; Harting et al. 1991a; Raczkowski and Rosenquist, 1983; Hoshino et al. 2010). The SC is a laminar structure commonly subdivided into two functional parts. The superficial layers of SC (sSC) form functional unit purely related to vision, while intermediate and deep layers are involved in multimodal sensory integration and the generation of motor commands related to saccadic eye- or head-movements towards an object of interest (reviewed in Moschovakis, 1996; Stuphorn et al. 2000; Munoz, 2002; Nagy et al. 2006). The main visual input to sSC originates in the retina (Graybiel, 1974, 1975; Hoffmann, 1973; for review see Waleszczyk et al. 2004; May, 2006), but the sSC are also the target of descending projections from multiple visual cortices (Harting et al. 1992).

There are a limited number of publications concerning the presence of oscillations and/or oscillatory synchronization of neuronal activity in the optic tectum (cat SC: Mandl, 1993; Brecht et al. 1998, 1999; 2001; Chabli et al. 2000; birds: Neuenschwander et al. 1999; Goddard et al. 2012; ferret: Stitt et al. 2013; monkey: Le et al. 2007; rat: Baranauskas et al. 2016). Moreover, these reports are somewhat inconclusive, in that some data showed oscillations in the SC that were dependent on stimulus velocity (Mandl, 1993; Chabli et al. 2000; Pauluis et al. 2001), while other data showed a lack of dependency on stimulus velocity (Brecht et al. 1998, 2001). In order to clarify this issue, we identified and characterized oscillations in the “spontaneous” (background) and in visually evoked activity in the cat’s sSC. We asked whether sSC oscillations were induced by visual stimulation or were inherited from the background activity and enhanced by visual stimulation. We also asked whether phase-locked to visual stimulus oscillations resulted from a visual stimulus-related phase reset of oscillations present in the background activity or whether these oscillations were generated by an independent mechanism.

Our results show that oscillations are common in background and stimulus-evoked activity of sSC neurons. Oscillations related to the visual stimulus can appear in two forms: oscillations that are stimulus-phase-locked to the stimulus and oscillations that are stimulus-phase-independent of the stimulus. In addition, we showed that phase-locked to stimulus oscillations are dependent on stimulus dynamics. Our results suggest that oscillations that are stimulus-phase-independent of the stimulus may be generated by the same mechanism as “spontaneous” oscillations, while oscillations that are stimulus-phase-locked to the stimulus may result from a phase reset of oscillations present the background activity or/and from intracollicular interactions.

Preliminary reports have been published in abstract form (Foik et al. 2010; 2011a, b).

## Materials and Methods

### Ethical Approval

Acute experiments were performed on anesthetized adult cats of either sex. All experimental procedures were carried out to minimize animal numbers and suffering, followed regulations and standards of the Directive 2010/63/EU of the European Parliament and of the Council of 22 September 2010 on the protection of animals used for scientific purposes. Procedures were approved by the First Warsaw Local Ethics Committee for Animal Experimentation (permission no.: 77/2010) and complied with the Polish Animal Protection Law.

### Surgical procedures

A typical experiment lasted four days, during which time neuronal activity from the sSC was recorded, apart from short breaks needed to change electrode tracks. Cats were given dexamethasone phosphate (0.3 mg kg^−1^, i.m.; Rapidexon, Eurovet Animal Health BV, Netherlands) to reduce brain edema on the day preceding the experiment. Animals were initially anesthetized with a mixture of xylazine (3 mg kg^−1^, i.m.; Xylavet, ScanVet), propionylpromazine (1 mg kg^−1^, i.m. Combelen, Bayer), and ketamine (20 mg kg^−1^, i.m.; Ketanest, BioVet). Tracheal and cephalic vein cannulations were performed to allow, respectively, artificial ventilation and infusion of paralyzing drugs. Bilateral sympathectomy was performed to further minimize eye movements. In the case of one cat, initial anesthesia was induced with midazolanium (0.05 ml kg^−1^, i.m., Midanium, WZF Polfa S.A., Poland), butorphanol (0.04 ml kg^−1^, i.m., Butomidor, WZF Polfa S.A, Poland) and medetomidine hydrochloride (0.04 ml kg^−1^, i.m., Domitor, WZF Polfa S.A., Poland). Initial anesthetic procedures had no effect on the obtained experimental results. Anesthesia was maintained with a gaseous mixture of N_2_O/O_2_ (2:1) and isoflurane (Baxter, Poland) of 1 – 1.5% during surgical procedures and 0.3 – 0.6% during recordings to keep monitored anaesthesia care value equal 1 (MAC = 1.0) and physiological parameters relatively constant (heart rate, blood saturation and presence of slow oscillations as theta and delta waves in the ECoG). Antibiotics (enrofloxacin, 5 mg kg^−1^, Baytril, Bayer Animal Health GmbH, Germany), dexamethasone phosphate (0.3 mg kg^−1^), and atropine sulfate (0.1 mg kg^−1^, to reduce mucous secretion, WZF Polfa S.A., Poland) were injected i.m. daily. Paralysis was induced with an i.v. injection of 20 mg gallamine triethiodide (Sigma - Aldrich) in 1 ml of sodium lactate solution and maintained by a continuous infusion of gallamine triethiodide (7.5 mg kg^−1^ hr^−1^, i.v.) in a mixture of equal parts of 5% glucose and sodium lactate. Body temperature was automatically maintained at ~ 37.5 °C with an electric heating blanket. Animals were artificially ventilated and expired CO_2_ was continuously monitored and maintained between 3.5 – 4.5% by adjusting the rate and/or stroke volume of the pulmonary pump. The heart rate and the electrocorticogram (ECoG; field potential recorded from the surface of the occipital lobe) from the side contralateral to the recording site were monitored continuously. A heart rate below 220 bpm and slow-wave synchronized cortical activity were maintained by adjusting the isoflurane level in the gaseous N2O and O_2_ mixture. Atropine sulfate (1-2 drops, 0.5% Atropinum Sulfuricum, WZF Polfa S.A, Poland) and phenylephrine hydrochloride (1-2 drops, 10% Neo-Synephrine, Ursapharm, Germany) were applied daily onto the corneas to dilate the pupils and retract the nictitating membranes. Air-permeable zero-power contact lenses were used to protect the corneas from drying. Lenses of other power were used if needed.

A fiber optic light source was used to back-project the optic discs onto a spherical screen located 0.7 m in front of the cat (Pettigrew et al. 1979). The positions of the *areae centrales* were plotted by reference to the optic discs (Bishop et al. 1962).

### Recording and visual stimulation

Extracellular single-unit recordings were made from neurons located in the *stratum griseum superficiale, stratum opticum,* and the upper part of the *stratum griseum intermediale* (collectively referred to as sSC). A plastic cylinder was mounted and glued around a craniotomy positioned above the SC (Horsley - Clarke coordinates P1 - A5 and L0 - L5). The cylinder was filled with 2% agar gel and sealed with warm wax. Action potentials were recorded with tungsten, stainless-steel (6-10 MΩ; FHC, Brunswick, ME), or platinum-iridium microelectrodes (1-2 MΩ; Thomas Recording GmbH, Germany), conventionally amplified, monitored *via* a loudspeaker, and visualized on an oscilloscope. Recorded signals were digitized and fed to a PC computer for online display, analysis, and data storage using CED 1401 Plus or Power 1401 mk II and Spike2 software (Cambridge Electronic Design, UK). Signals containing spike waveforms were band-pass filtered between 0.5 - 5 kHz and digitized at a sampling rate of 50 kHz. The ECoG was band-pass filtered between 0.1 - 100 Hz and digitized at a sampling rate of 1 kHz. Neuronal responsiveness to visual stimulation and the eye of origin (ipsi- and/or contralateral) were determined and the excitatory receptive fields (RFs; minimum discharge fields) plotted using black or white hand-held stimuli. Ocular dominance was determined by listening to the neuronal responses *via* a loudspeaker (spikes converted to standard TTL pulses). The dominant eye was chosen for subsequent visual stimulation (with the other eye covered). If conditions allowed, i.e. the registered signal was stable with well-isolated single unit activity, responses were also recorded during stimulation of the non-dominant eye.

To reduce the risk that observed oscillations, in fact, reflect phase locking to the refresh rate of the monitor (Butts et al., 2007; Williams et al., 2004; Wollman and Palmer, 1995; Koepsell et al., 2009) we used a slide projector under computer control for visual stimuli presentation. We recorded spontaneous activity and responses to a light rectangle (1 x 2 deg or 0.5 x 1 deg; 4 - 6 cd (m^2^)^−1^ stimulus luminance against a 0.5 - 1 cd (m^2^)^−1^ background luminance) flashing (0.5 – 1 s stimulus onset and offset) or moving with a constant velocity ranging from 2 to 1000 deg s^−1^ (velocity values were approximately uniformly distributed on a logarithmic scale). Stimuli were projected onto a concave spherical screen located 0.75 m in front of the cat and covering a 70 deg area of the visual field. The center of the screen was adjusted to overlap the recorded RF center, and a constant velocity stimulus moved through the RF center along its horizontal or vertical axis. Stimulus movement was achieved by computer-controlled rotation of a mirror attached to the axle of a galvanometer. Voltage changes transferred to the galvanometer were generated by Spike2 software and the digital-analog converter, CED 1401 Plus or Power 1401 mk II. To ensure smooth stimulus movement, a single sweep with a full 50 deg amplitude was achieved in 500 voltage steps for the fastest stimulus (1000 deg s^−1^) and up to 5000 steps for the slowest (2 deg s^−1^). Each trial consisted of motion in one direction followed by a 1 s pause and then motion in the reverse direction at the same velocity followed by 1 s pause. The number of trials was roughly proportional to stimulus velocity (10 repetitions for 2 deg s^−1^ up to 100 repetitions for 1000 deg s^−1^).

The RF spatiotemporal profiles (or maps) were plotted according to the averaged responses to a flashing light rectangle pseudo-randomly presented at 25 locations along the horizontal or vertical axis of the RF. Recordings for stimulus ON and OFF responses lasted 0.5, 0.8 or 1 s.

Only records that contained data without visual stimulation for more than 60 s were used for background (spontaneous) activity analysis.

### Localization of recording sites

Small electrolytic lesions (20 μA, 10 s) were made at the end of recording sessions. During some experiments, electrodes were coated with DiI (1,1’-dioctadecyl-3.3,3’,3’-tetramethylindocarbocyanine perchlorate; Sigma-Aldrich, Germany) to facilitate electrode tract reconstruction (DiCarlo et al. 1996). Animals were killed by an overdose of sodium pentobarbitone (Nembutal^®^ Sodium Solution, Abbott Laboratories, i.v.), the brains removed and then immersed in 4% paraformaldehyde in 0.1 M phosphate buffer (pH 7.4). Electrode tracks were reconstructed from 50 μm coronal slices stained with cresyl violet; the analysis confirmed that recordings were obtained from the sSC.

### Data analysis

Spike2 (Cambridge Electronic Design, UK) software was used for spike sorting and offline conversion of single unit waveforms into discrete spike time occurrences. Spike sorting was based on waveform analysis. We analyzed 122 recorded sSC units out of a total of 168, for which we were sure that every spike was correctly classified during offline sorting. In all cases, single unit discrimination was confirmed by the presence of an absolute refractory period in the interspike interval histogram (ISI; e.g. Fig. *1D*). Two units that recorded simultaneously were discriminated only when the shape of the two spike templates differed substantially (e.g. Fig. 1*B, C*). Further data analyses were performed using Matlab software (The MathWorks Inc., Natic, MA).

**Figure 1.**
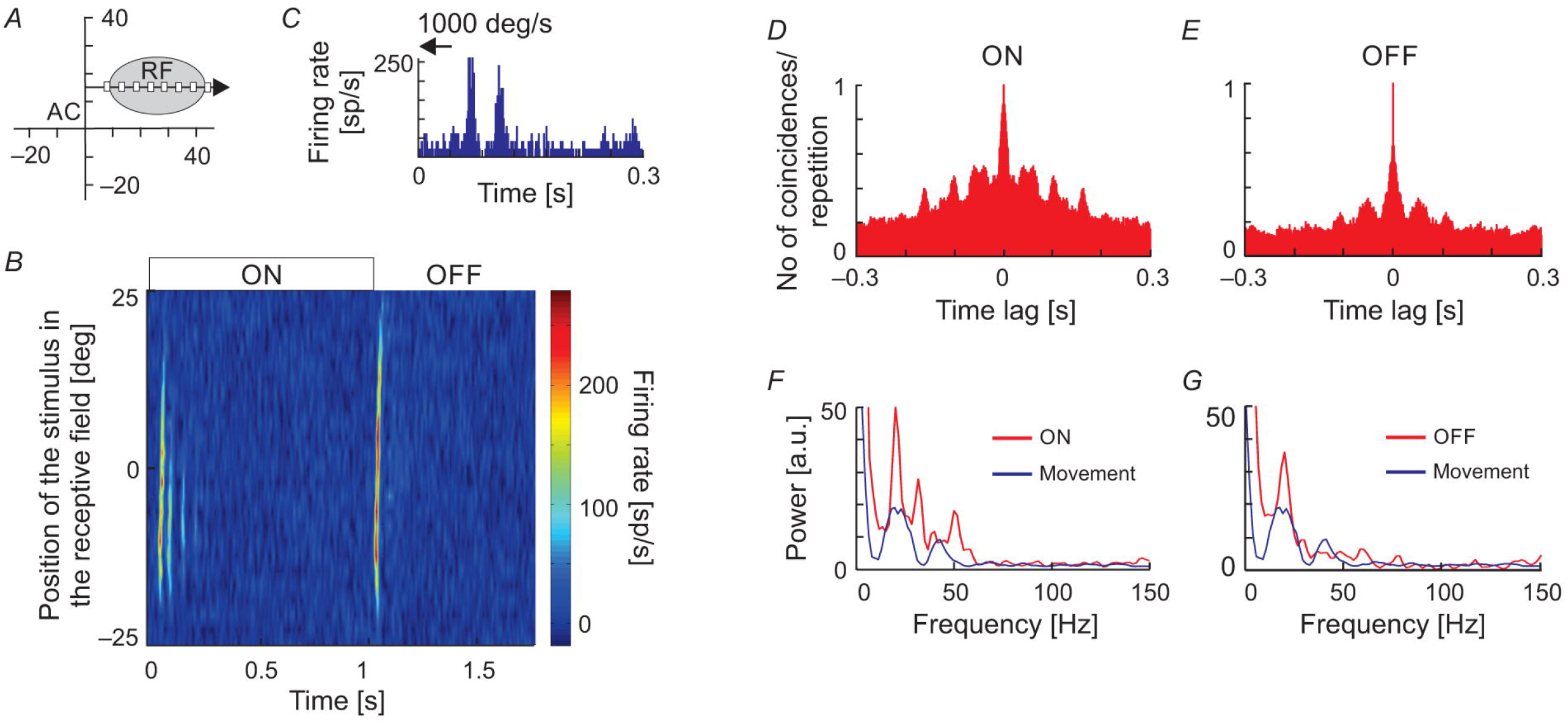
Extracellular neuronal recordings of oscillatory activity from the superficial layers of the SC. (A), Top panel shows the raw signal recorded in response to a light bar moving at a velocity of 5 deg s^−1^ back and forth through the receptive field (RF). In this and all subsequent figures, horizontal arrows indicate the onset, duration, and direction of the moving stimulus. The bottom panel represents an enlarged part of the raw signal. An oscillatory pattern is clearly visible as single spikes or doublets appearing with a constant interval. (B), Spike waveforms of two cells selected from the recording in (A). Only the unit with the higher amplitude (black line) was considered for further analysis. (C), Spike waveforms classified into two putative units based on the principal component analysis. Colors mimic those of the waveforms in panel (B). (D), Inter-spike interval (ISI) histogram for the unit in (A) and (B) (black waveforms). The first peak reflects bursting activity (seen here as doublets), the other peaks indicate oscillatory activity. Insert represents an enlarged view of the beginning of the ISI to emphasize the presence of the refractory period. (E), Peri-stimulus time histogram (PSTH) divided into 200 bins for 10 repetitions of the stimulus presentation for the unit in (A-D). (F), Corrected autocorrelogram (cACH) from the signal in (A); resolution 0.5 ms. (G), Power spectrum calculated from the cACH in (F).

Data analyses were based on sets of spike trains of background activity (lasting at least 60 s), spike trains corresponding to repetitive stimulus motion, or records obtained with the flashing spot, as described above. Peri-stimulus time histograms (PSTHs) were constructed based on the responses to all repetitions of a given stimulus; PSTH bin length was always 0.5 ms. In the following figures, the PSTH time bases were usually divided into 200 bins regardless of the single trial duration for ease of presentation. This approach resulted in different bin lengths that were dependent on stimulus velocity, but in no case did this change the characteristics of the presented response.

#### Autocorrelation and Fast Fourier Transform

Raw autocorrelation histograms (rACHs) were computed following the method introduced by Perkel and co-workers (1967a, b; see also Aertsen et al. 1989) and with normalization similar to that proposed by Bair and co-workers (2001) and Kohn and Smith (2005). The rACH computed for single spike trains was assessed according to the following equation:

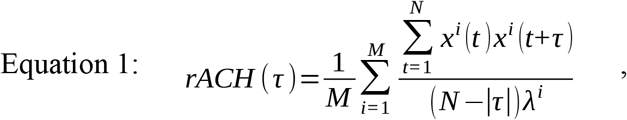

where *M* is the number of trials, *N* is the number of bins in the trial (the same in all trials), *i* is a given trial, *τ* - a time lag in the number of bins, *λ* is the mean spike count in the *i^th^* trial, and *X^i^(t)* is the spike train related to the *i^th^* trial.

In the case of an averaged PSTH, *M* = 1, thus ACH for a PSTH (ACH_PSTH_) was assessed according to the following simplification of equation 1:

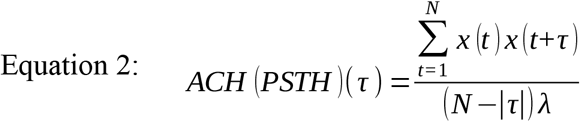

where *N* is the number of PSTH bins, *x(t)* is the spike count in bin t of the PSTH, *τ* - particular time lag in the number of bins, and *λ* is the mean spike count per bin averaged over the whole PSTH.

rACHs were prepared from single spike trains or from PSTHs divided into 0.5 ms bins with a maximum lag of 0.3 s. To obtain corrected ACH (cACH), a shift predictor (Equation 3) was computed and subtracted from the rACH to remove stimulus influences:

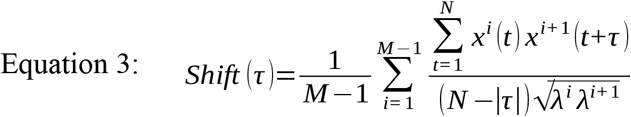

The temporal dynamics of ACHs (raw and corrected) during stimulus presentation were assessed using a sliding-window method (ACHs in time). A 250 ms window (500 bins of 0.5 ms duration) was shifted along the time axis with resolution dependent on stimulus velocity; 10 ms for stimulus velocities of 20, 50, 100, 200, 500, and 1000 deg s^−1^; 20 ms for 10 and 5 deg s^−1^; and 25 ms for 2 deg s^−1^. For ACHs in time, the maximum time lag was 0.25 s with a 0.5 ms bin width.

Following the rules adopted by Samonds and Bonds (2005), a cell was classified as oscillatory when its ACH contained a second oscillatory peak (e.g. Fig. 1*F*). Moreover, the amplitude of a prominent peak in the power spectrum computed from the ACH had to exceed at least twice the amplitude of fluctuations assessed by z-statistics (see also section: *Estimation of oscillation strength*).

To identify oscillation frequencies, amplitude spectra were computed from both raw and corrected ACHs using a MatLab algorithm for Fast Fourier Transform (FFT) without any window. For ACHs in time, an FFT was computed for each time window in each repetition, averaged over the repetitions, and then presented in the form of a twodimensional matrix showing the temporal dynamics of the amplitude spectra.

#### Estimation of oscillation strength

The strength of oscillation index (SO) was estimated by a modified z-score standardization method (Wypych et al. 2012). The SO was defined as the difference between the amplitude of the prominent peak (*F*) in the power spectrum and the mean of the power spectrum (*mean* (FFT)) computed from the ACH (this is in contrast to the power spectrum computed from a raw signal used by Wypych and colleagues (2012)), and divided by the standard deviation of power spectrum (*std*(FFT)):

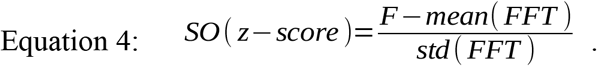

Thus, the SO(z-score) represents the distance between the peak amplitude in the power spectrum and the mean power spectrum in units of the standard deviation. As a result of such standardization, the index is resistant to the number of coincidences in ACHs. As has been shown earlier, estimation of the SO from the spike train with a z-score normalization allows for the detection of weak oscillations in cases of low spiking activity (Wypych et al. 2012). However, in such cases, the index is dependent on the length of the recording period. Here we calculated the SO(z-score) from ACHs of equal length, which makes the index essentially independent of spike train length. Additionally, the strength of oscillation was also estimated using the *oscillation score* (Os) introduced by Mureşan and colleagues (2007):

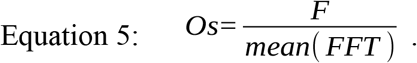

Both measures, SO(z-score) and Os, are dimensionless. SO(z-score) and Os were assessed from the mean and standard deviation over a frequency range of 8 – 100 Hz for visually evoked activity and from 5 to 100 Hz for background activity.

#### Renewal density analysis

Renewal density analysis (RDA) (Mountcastle et al. 1990; Lebedev and Wise, 2000; Samonds and Bonds, 2005) was performed to reveal the source of oscillatory activity in sSC cell (intrinsic or extrinsic). Inter-spike intervals were randomly shuffled and surrogate (shuffled) spike trains were reconstructed to destroy the temporal order of the intervals. Surrogate data obtained by this procedure were further analyzed in the same way as the original (non-shuffled) experimental data. We compared the spectra of different types of ACHs (raw and corrected) prepared for shuffled and original data, and the amplitudes at the dominant frequency for background and stimulus-evoked oscillations. Shuffling of the interspike intervals and subsequent analysis of spike trains obtained for visual stimulation were limited to the period of the visual response (duration of visual stimulation plus 300 ms following stimulation). Any amplitude decrease at the dominant frequency was considered to be an indication of an extrinsic source of oscillations (see Samonds and Bonds, 2005).

#### Statistical analyses

All statistical analyses were performed using the Matlab software. Significant differences between the two populations were examined using a two-tailed Wilcoxon signed-rank test for related data and a Mann–Whitney U-test for unrelated samples. The mean and standard deviation (mean (SD)) are given to characterize the sample. Statistical differences between two distributions were revealed by a Kologomorov-Smirnov (K-S) test. The relationship between the two variables was assessed with the Pearson correlation coefficient (*r*). Differences and correlations were considered significant for *p* ≤ 0.05.

## Results

The results are based on spike train analysis from 122 SC cells and primarily responses to moving stimuli. The only limited analysis is given for flashing spot stimuli. Visually responsive cells were recorded from superficial layers of SC: *stratum griseum superficiale*, *stratum opticum* and upper part of *stratum griseum intermediale*, as confirmed by electrode tract reconstruction. Oscillations were identified in the spontaneous activity of 58% (23 / 40) of sSC cells, and in the visually evoked activity of 41% (50 / 122) of sSC cells. We observed oscillations in an evoked activity that were phase-locked to the stimulus (i.e. stimulus on-set as in the case of moving or flashing spots or off-set as in the case of flashing spots) and oscillations that were stimulus-phase-independent but were generated or enhanced by visual stimulation. Stimulus-phase-locked oscillations were revealed by 1) the presence of apparent changes in the firing rate during and/or shortly after visual stimulation, that were clearly visible in the form of vertical stripes in the raster plots and as a specific temporal pattern in averaged PSTHs, 2) the presence of a second oscillatory peak in the ACH_PSTH_ computed from averaged PSTHs, as well as 3) the presence of a prominent peak (z-score > 2) in the computed from ACHPSTH power spectrum. Stimulus-phase-independent oscillations were identified by 1) the presence of a second oscillatory peak in the rACH computed from a spike train obtained for each single stimulus repetition and averaged over number of repetitions, and corrected by subtraction of the stimulus-dependent shift predictor, and 2) the presence of a prominent peak (z-score > 2) in the power spectrum computed from cACH. Since the phase of these oscillations varied in subsequent stimulus repetitions, they were not apparent in averaged PSTHs.

Figure 1 shows an example of oscillations in the visually evoked activity of sSC neuron. Oscillations are clearly visible as regularly encountered spikes (indicated by arrows in Fig. 1*A*) in the raw signal recorded in response to a light bar moving (5 deg s^−1^) back and forth along the horizontal axis of the RF. Strong oscillatory activity (stimulus-phase-independent) is present despite the weak response to the stimulus (shown in Fig. 1*E*). Due to various phase of oscillations relative to visual stimulation these oscillations are not apparent in PSTH (Fig. 1*E*), however they could be easily identified in the rACH and cACH (Fig. 1*F*) as a periodic increase in the number of coincidences in a interval that is equal to the intervals between doublets of spikes and/or single action potentials (see Fig. 1*A*). A similar temporal interval can be deduced from the oscillatory peaks in the cACH (Fig. 1*F* and in the ISI (indicated by the arrow in Fig. 1*D*).

Oscillatory frequency (28 Hz) determined from a dominant peak in the power spectrum (Fig. 1*G*), corresponding to a time lag of 0.036 s between single spikes or bursts of spikes in the raw signal, cACH, and ISI.

### Oscillations in background activity

Oscillations occurred in the background activity in 58% (23 / 40) of the sSC cells. Figure 2 shows examples of ACHs (Fig. 2*A, B*) and corresponding amplitude spectra (Fig. 2*C, D*) computed from the background activity for two sSC cells. Oscillatory frequencies in background activity differed between these SC neurons. In the whole analyzed sample, the frequencies of the spontaneous oscillations ranged from 5 to 90 Hz (Figure 2E). Oscillations in the spontaneous activity were not a stable phenomenon, that is, they were not always visible in every background activity registration of a given neuron. Although the strength of the spontaneous oscillations could differ between recordings, the oscillatory frequencies were very similar for a given cell whenever oscillations were detected with a mean oscillation strength (OS(z-score)) of 3.24 ± 0.95 SD and average frequency 38.8 Hz ± 30.42 Hz SD (Figure 2*F-G*).

**Figure 2.**
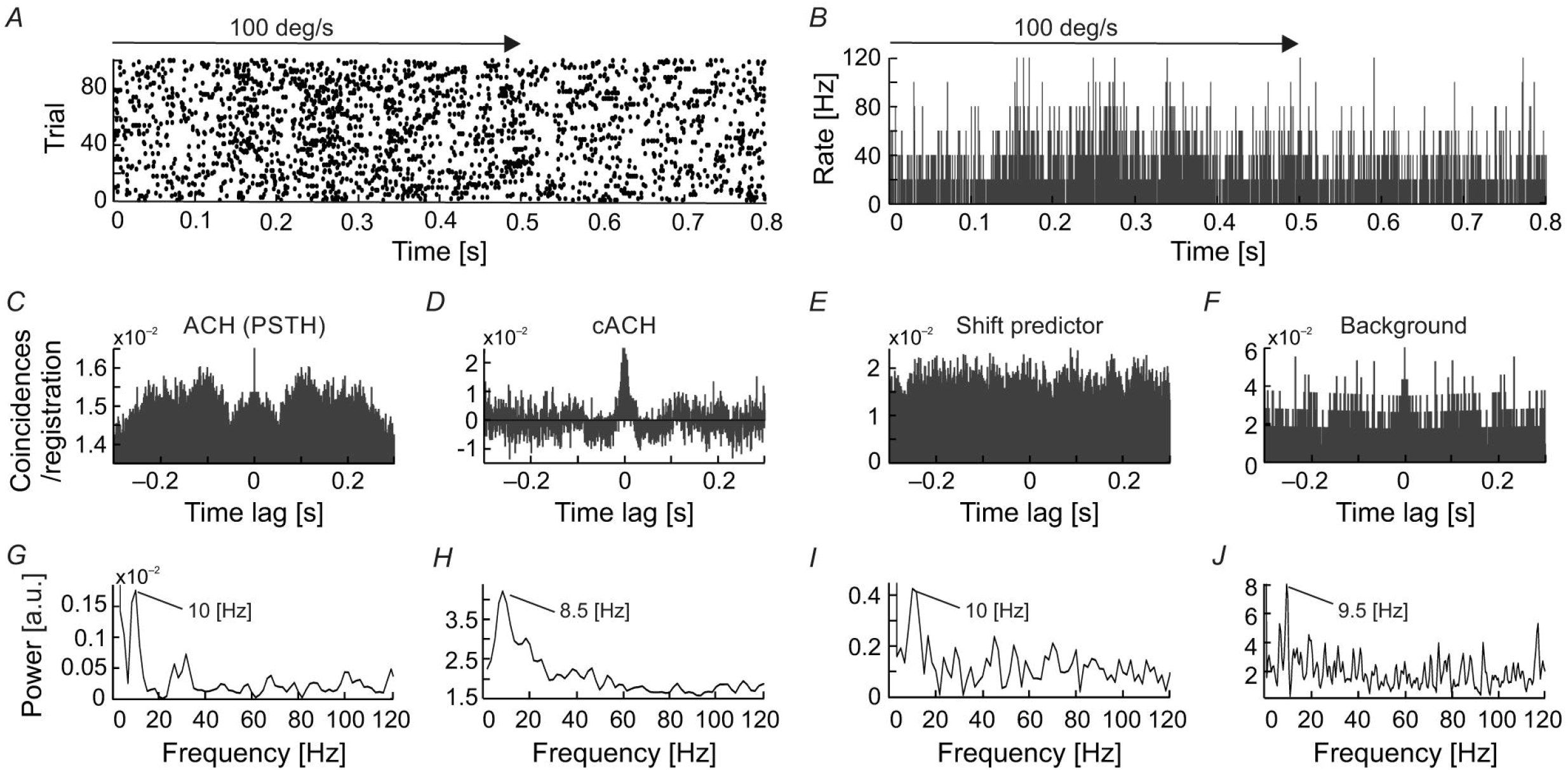
Examples of oscillations in the background activity of sSC neurons. (A), The raw autocorrelogram consisted of 1720 spikes of one of the sSC neurons, recorded over 262 s of spontaneous activity. (B), The autocorrelogram for the other sSC neuron consisted of 8320 spikes, recorded over 104 s. (C), (D), The respective power spectra computed from autocorrelograms shown in A and B. Note the different oscillatory frequencies for the two cells, 23 and 70 Hz. (E), Distribution of the frequencies of oscillations found in the background activity of the SC. (F), the median oscillation strength for background oscillations. (G), the median frequency of background oscillations. Bottom and upper boxes lines indicate 25^th^ and 75^th^ percentile. The error bars indicate minim and maxim values in the datasets.

### Visually evoked activity: oscillations that are stimulus-phase-independent of a moving stimulus

An example of stimulus-phase-independent oscillations is presented in Figure 1 (described above). To reveal oscillations that were not phase-locked to the stimulus onset, we analyzed single trials (rACH) and ACHs corrected after subtracting a shift predictor (cACH). Stimulus-phase-independent oscillations over a frequency range of 8 – 100 Hz were identified in 25% (31 / 122) of cells. A subset of these cells (14 / 31) also had recordings that allowed us to observe oscillations in the background activity, i.e. a period without visual stimulation lasting at least 60 s. For the majority of these cells (12 / 14) the frequency of stimulus-phase-independent and background oscillations were similar.

Figure 3 shows an example of a time-resolved ACH in response to a light bar moving at 1000 deg s^−1^ back and forth along the horizontal axis of the RF (PSTH in Fig. 3*A*). An increase in the number of coincidences in the ACH corresponded to an increase in the firing rate in response to stimulus movement (Fig. 3*B*). Amplitude spectra (Fig. 3*C*) computed from corrected ACHs revealed an increase in the strength of oscillations during the responses compared with periods of activity between stimulus movements, in pauses. A different frequency pattern and power spectrum were obtained for the forward and backward directions of stimulus motion.

**Figure 3.**
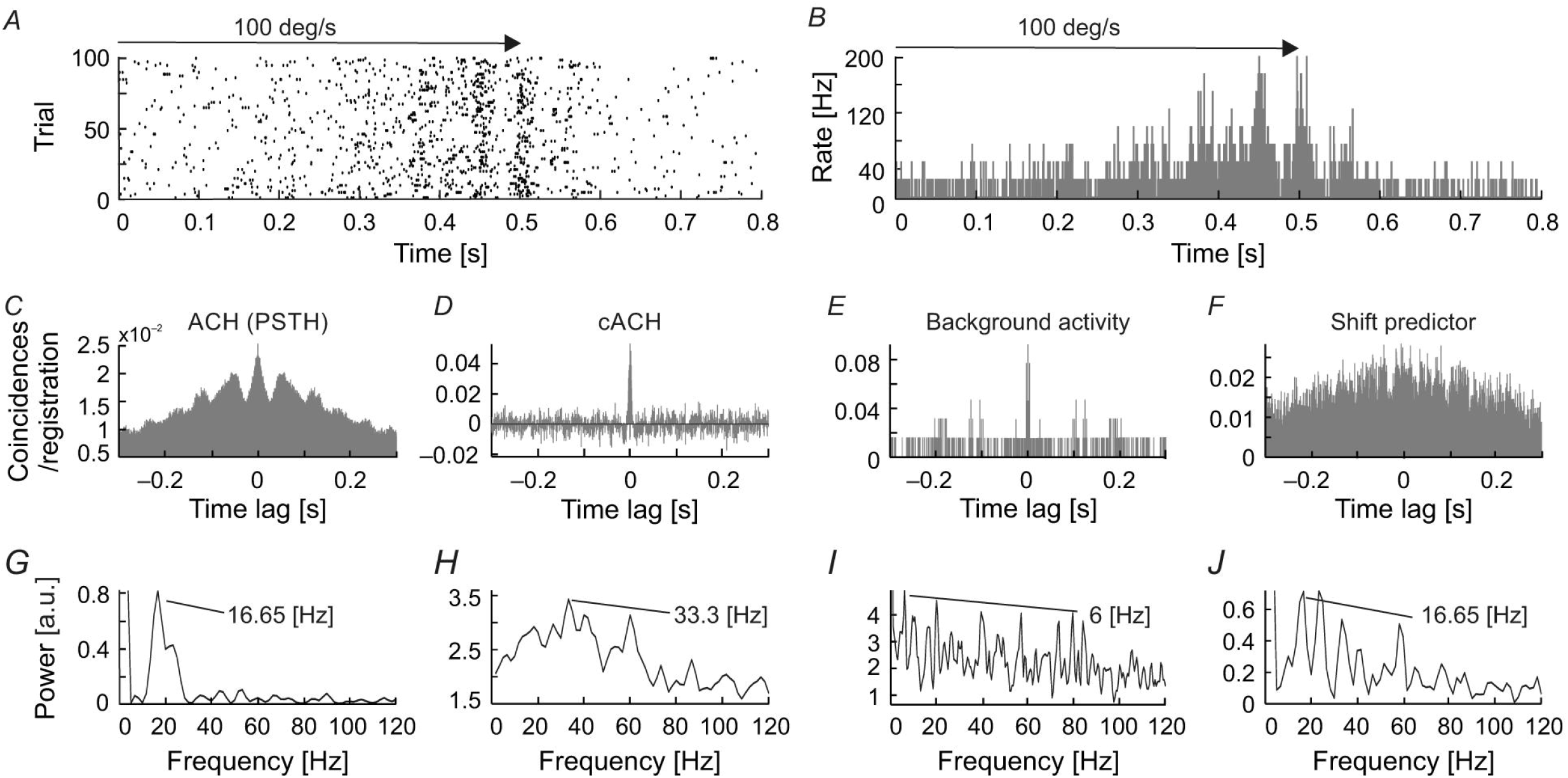
Time-resolved ACH showing oscillatory responses of an sSC neuron that responded with a different pattern of oscillatory frequencies for each direction of stimulus movement. (A), PSTH of the response to a light bar moving at 1000 deg s^−1^, averaged over 100 repetitions and divided into 200 bins. Small horizontal arrows above PSTH indicate the time and direction of stimulus movement. (B), Time-resolved raw ACH averaged over trials. A single ACH, shown to the left in red, corresponds to the maximum response for stimulus movement. Single ACHs were computed with a 0.5 ms resolution using a 250 ms sliding window with 10 ms shift for every stimulus repetition and then averaged over the repetitions. Indicated by arrows between (A) and (B) are ACHs computed for two first neighboring windows. The increase in the number of coincidences, coded by color, corresponds to an increase in the firing rate in the response shown in (A). (C), Spectrogram obtained by computing the power spectra (by subtracting a shift predictor) on corrected single ACHs shows the increase in the strength of oscillations at times corresponding to the responses shown in (A). Scale bars for the time-resolved ACH and spectrogram are shown on the right.

Similar stimulus-phase-independent oscillation frequencies were observed over a broad range of stimulus velocities (Fig. 4*A - C*). Figure 4 shows examples of amplitude spectra computed from time-resolved cACHs in response to different stimulus velocities (10, 50 and 1000 deg s^−1^) for one SC cell. The same dominant oscillatory frequency (33 Hz) is clearly visible in response to stimulus movement from right to left in all three plots of the power spectrum (red line in Fig. 4*D - F*) and was also present in the background activity (power spectrum plotted as blue line in Fig. 4*D - F*).

**Figure 4.**
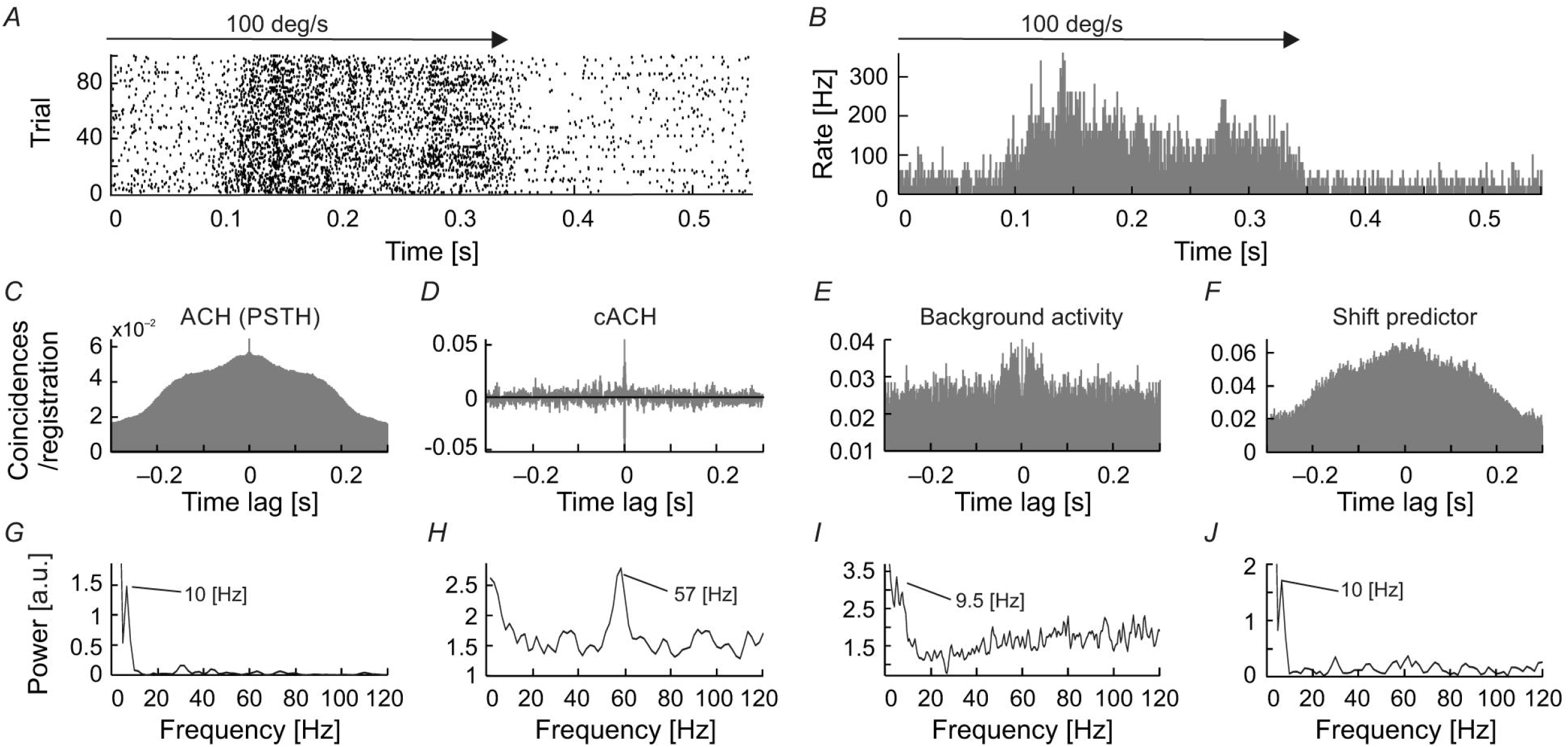
Responses of one sSC neuron to moving stimuli of different velocities (20, 100, and 1000 deg s^−1^). (A - C), PSTHs of neuronal responses for three velocities). (D – F), amplitude spectra (spectrograms) computed from ACHs for responses above. Power spectra were computed from corrected ACHs after subtracting shift predictors. Changes in the power spectra are coded according to the color bars on the bottom. The right-hand side of the panel shows a single example of the power spectrum (red lines) taken from the region representing the maximum power during the response. Amplitude spectrum computed for the background activity for the same neuron is shown as the blue line. Note the similar pattern of oscillation frequencies for visual stimulation and background activity.

### Visually evoked activity: oscillations that are moving stimulus-phase-locked

Stimulus-phase-locked oscillations, that is, oscillations occurring at the same time relative to the onset of visual stimulation and with the same phase for each stimulus repetition (for review see Koepsel et al. 2010), were found in the evoked activity of 16% (20 / 122) of the neurons tested with moving stimuli.

Figure 5 shows examples of the occurrence of stimulus-phase-locked oscillations in response to stimulus movement away from the vertical meridian of the visual field along the horizontal axis of the RF (Fig. 5). To assess the frequency and strength of stimulus-phase-locked oscillations, power spectra were obtained from ACHPSTHs computed from PSTHs averaged over all stimulus repetitions (Fig. 5*A-L*). The strength of the stimulus-phase-locked oscillations clearly depended on stimulus velocity and was greater for faster stimulus velocities (see a representative example of the cell in Fig. 5*M*). Oscillations of this type were rarely observed at very low velocities (below 20 deg s^−1^). In the majority of cells (15 / 20) a typical pattern of stimulus-phase-locked oscillations, apparent in the form of vertical stripes in the raster plots, emerged at 50 deg s^−1^ (e.g. Fig. 5*B, E, H, K*), and were clearly visible in responses to stimuli moving at higher velocities (e.g. Fig. 5*C, F, I, L*). In a smaller subset of cells (5 / 20) this characteristic pattern was already apparent with stimulus velocities of 20 deg s^−1^. We have observed differences in the strength of stimulus-phase-locked oscillations of the same frequency for opposite directions of movement. A typical example of this is shown in Figure 6*A – D*, in which stronger oscillations are seen for stimulus movement from right to left (Figure 6*E, F*). Suppression of the firing rate at the beginning of the response is visible in both Figures 5*C* and 6 *A, B* and was a characteristic feature in the majority of recordings showing stimulus-phase-locked oscillations. Figure 6*G* shows the distribution of frequency differences between oscillations found for two distinct directions of movement. More than half of cells (54%) showed the difference in frequencies in responses to different directions. The scatter plot in Figure 6*H* shows the greater strength of oscillations in a direction away from *area centralis* (in our case from left to right).

**Figure 5.**
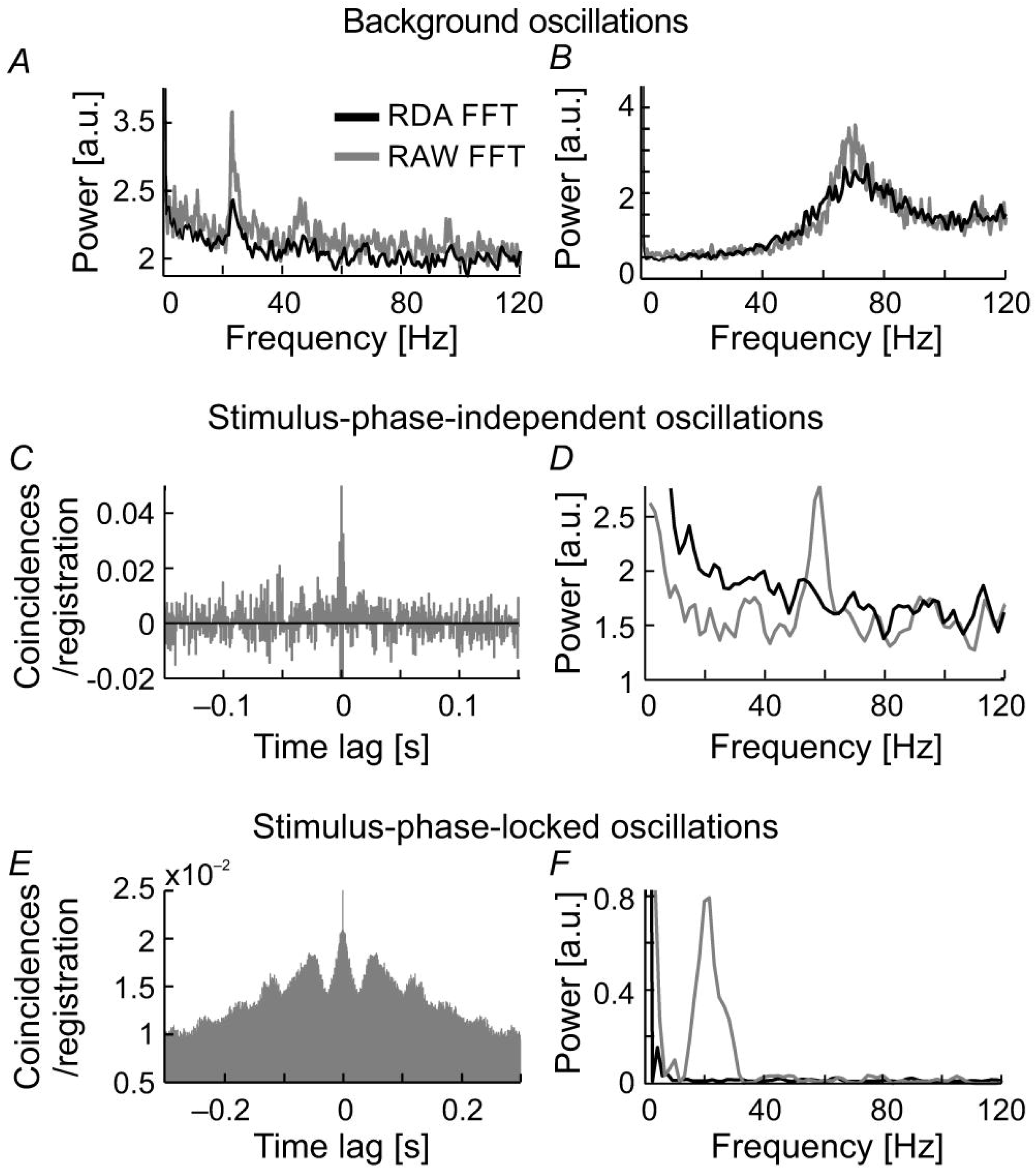
Changes in the temporal pattern of spikes generated by an SC neuron in response to moving stimuli at different velocities and in the same direction. (A - C), Raster plots for three stimulus velocities: 10, 50, and 200 deg s^−1^ respectively. (D - F), Corresponding PSTHs (100 bins). G, H, I, ACHs computed from the PSTH reveal the presence of stimulus-phase-locked oscillations for velocities above 10 deg s^−1^. (J - L), Amplitude spectra prepared from (G), (H), (I). (M), Plot of the strength of oscillation index (SO) as a function of stimulus velocity showing a positive correlation between the SO and stimulus velocity for this neuron. (N), The position of the RF in the visual field. Stimulus motion (indicated by the arrow) was directed away from the *area centralis* (AC). Note the changing temporal structure and increasing synchronization the of the responses in the raster plots and PSTHs as stimulus velocity increases, which is confirmed in the power spectrum analysis and is associated with a decrease in the firing rate at the beginning of the response, clearly visible for 200 deg s^−1^ in plot (C) and (F).

**Figure 6.**
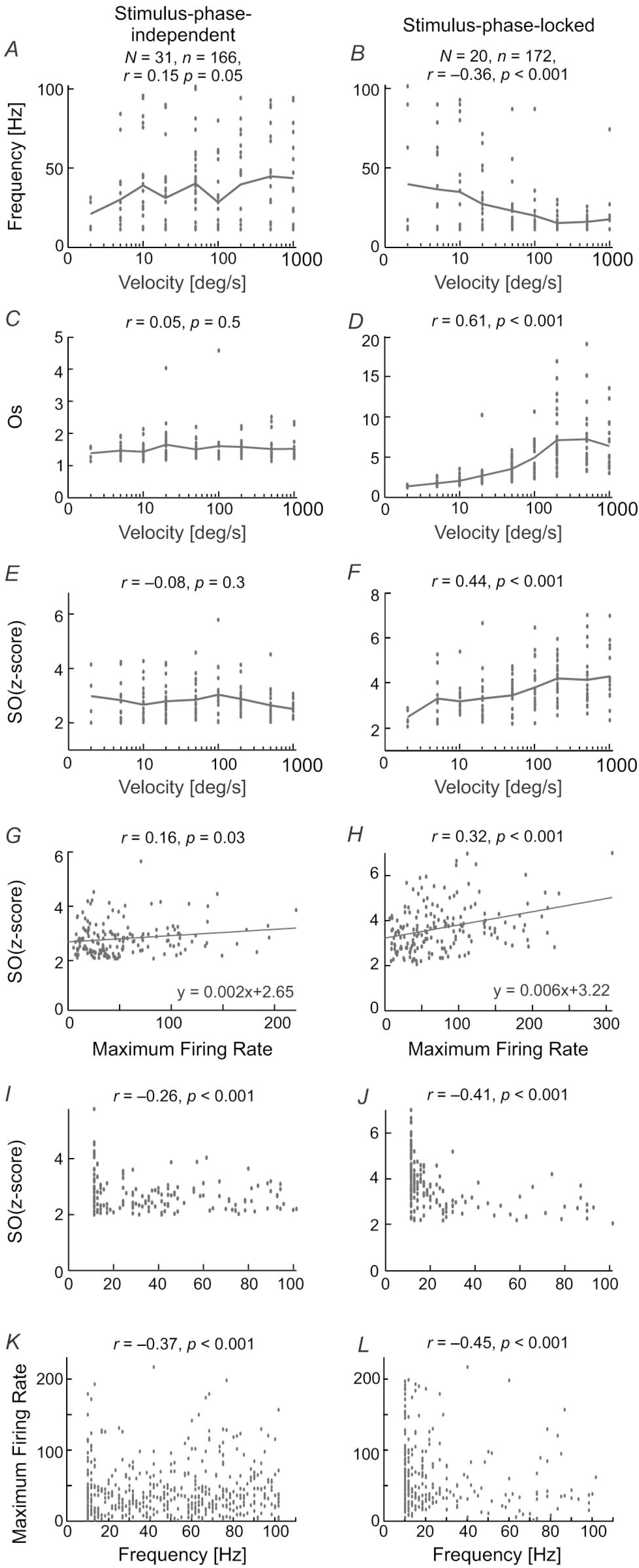
Different patterns of responses for a visual neuron in the SC evoked by reversing stimulus direction. (A), (B), Raster plots showing responses to visual stimuli moving to opposite directions at 200 deg s^−1^. (C), (D), PSTHs for responses shown in (A) and (B), bin size 0.5 ms. (E), (F), Amplitude spectra of the corresponding ACHs_PSTH_ computed from (C) and (D). Prominent peaks show the dominant frequencies in the responses. Note the clear decrease in firing rate at the beginning of the response for both directions in the raster plots. (G), the distribution of differences between frequencies found during responses to two directions of stimulus movement. (H), the scatter plot showing a shift of oscillation strength to the direction of movement from left to right. The black solid line indicates a linear fit to the data.

### Visually evoked activity: oscillations that are stationary stimulus-phase-locked

Temporal response patterns in the form of vertical stripes in the raster plots, typical for stimulus-phase-locked oscillations evoked by moving stimuli, could result from inhomogeneity in the spatial structure of the RFs. To exclude such a possibility, we recorded neuronal responses to flashing stimuli presented at random locations along the axis of stimulus movement within the RF (Fig. 7*A*) and constructed spatiotemporal RFs. Although maps of spatiotemporal RFs revealed multiple ON and OFF subregions, in 36% of cases (18/50) they also revealed the presence of, particularly strong oscillations in the ON responses. Figure 7*B* shows an example of a spatiotemporal RF of an SC neuron exhibiting strong stimulus-phase-locked oscillations in responses to fast stimulus movement (Fig 7*C*). In this example, strong oscillatory responses to stimulus ON-set were present and usually limited to the RF center. Although not apparent in the plot of the spatiotemporal RF (Fig. 7*B*), the stimulus-phase-locked oscillations were also present in responses to stimulus OFF-set as revealed in the ACHPSTH and its power spectrum (Fig. 7*E* and *G* respectively). Comparison of power spectra computed from ACHS_PSTH_ obtained for responses to stationary and moving stimuli (Fig. 7*F* and *G* red and blue lines, respectively) revealed the same dominant frequency of the stimulus-phase-locked oscillations in both cases. The similarity between frequencies of stimulus-phase-locked oscillations in response to moving and flashing stimuli were also observed for the other 17 neurons studied. Thus, we conclude that despite the inhomogeneities in the spatial structure of SC RFs, the typical pattern of stimulus-phase-locked oscillations in response to moving stimuli was related to the temporal structure of the spike trains and not due to the spatial structure of the RF.

**Figure 7.**
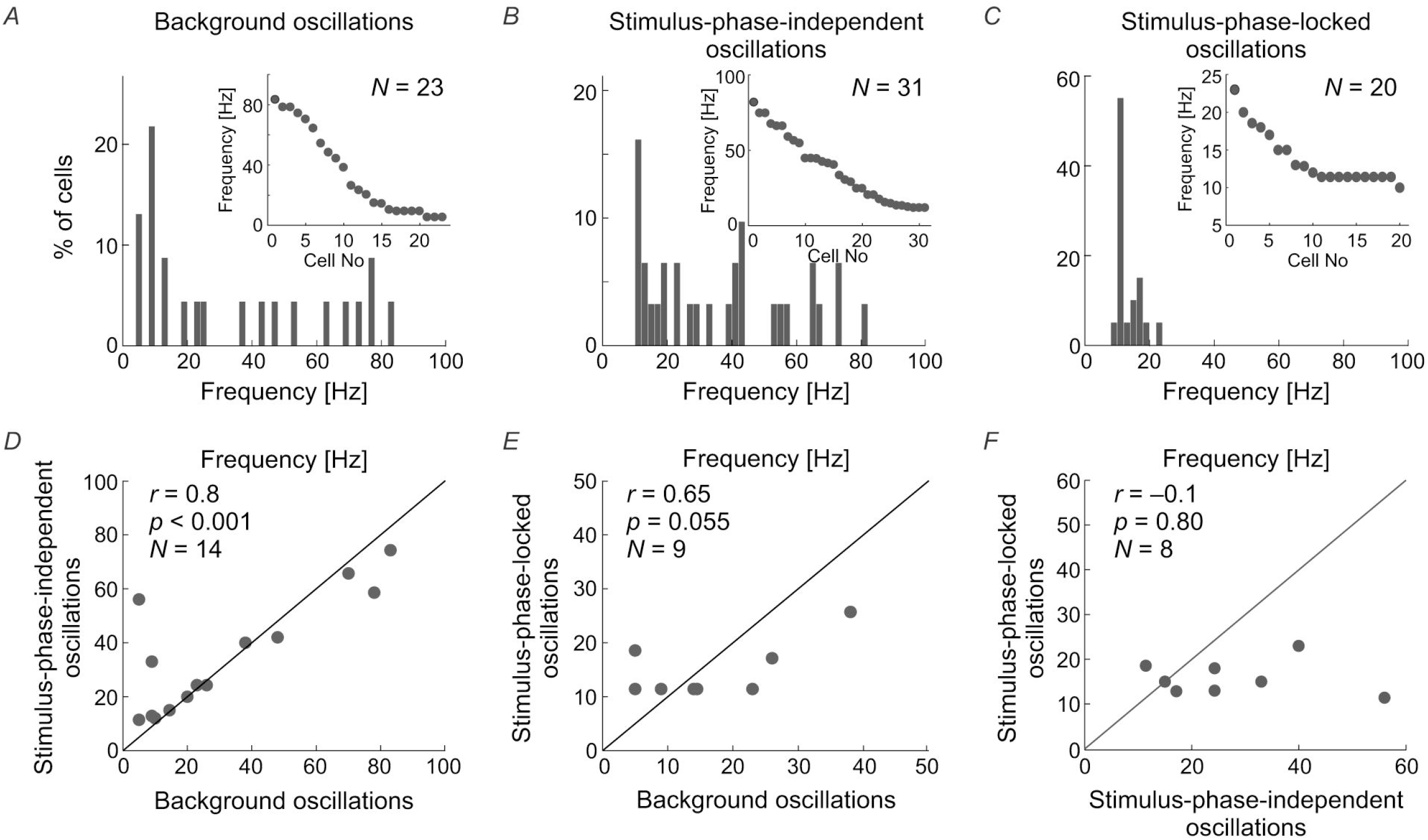
Oscillations in the responses of SC neuron to the ON-set, OFF-set, and movement of the light spot. (A), The position of the RF in the visual field and the scheme of visual stimulation with a moving or flashing spot (white rectangle). The black arrow indicates axis and direction of stimulus movement. (B), The spatiotemporal RF map shows oscillatory responses to the ON-set and OFF-set of the stimulus in multiple locations of the RF. Background activity was subtracted for better visualization of the stimulus-phase-locked oscillations. (C), Oscillatory response to fast movement (1000 deg s^−1^) across the RF. (D), (E), ACHs for ON and OFF responses in the RF location of strongest appearance of oscillations following stimulus ON-set, respectively. (F), (G), Amplitude spectra from (D) and (E) (red lines), respectively. The blue lines correspond to the amplitude spectrum computed from the ACHs in response to a moving stimulus. Note that the frequency of oscillations found in response to flashing spot (ON and OFF responses in (B)) in the RF is similar to that found in response to stimulus movement. The data excludes the structure of RF as the origin of stimulus-phase-locked oscillations.

### Simultaneous presence of more than one type of oscillation

To directly compare frequencies of different types of oscillations we have searched for the simultaneous presence of more than one type of oscillatory activity for a single neuron. Further, we have plotted raster plots and PSTHs to show the presence of oscillations in the raw data visible as fluctuations in firing pattern. Such fluctuations are also presented in various autocorrelograms and corresponding amplitude spectra to reveal and compare dominant frequencies of oscillations. The simultaneous presence of similar frequencies in the background oscillations and in the cACH may suggest enhancement of background oscillations by visual stimulation. The presence of similar frequencies in the background activity and in the ACH_PSTH_ as well as in the shift predictor may suggest that stimulus causes not only increase in strength of oscillations but also their locking and synchronizing to stimulus onset. In eight cells (7%; 8 / 122) we observed the simultaneous presence of two types of oscillations, a stimulus-phase-independent and stimulus-phase-locked (e.g. Fig. 8). For seven of these neurons, we also observed oscillations in background activity. In all these cells, the background and stimulus-phase-independent oscillations had similar frequencies. In the example shown in Figure 8, the frequency of the background oscillations (Fig. 8*F, J*) was similar to the stimulus-phase-independent (Fig. 8*D, H*) and stimulus-phase-locked oscillatory frequency (Fig. 8*C, G*) which confirms the hypothesis about enhancement and phase reset of background oscillations by the stimulus. A common dominant frequency around 10 Hz was also visible in the shift predictor (Fig. 8*E, I*).

**Figure 8.**
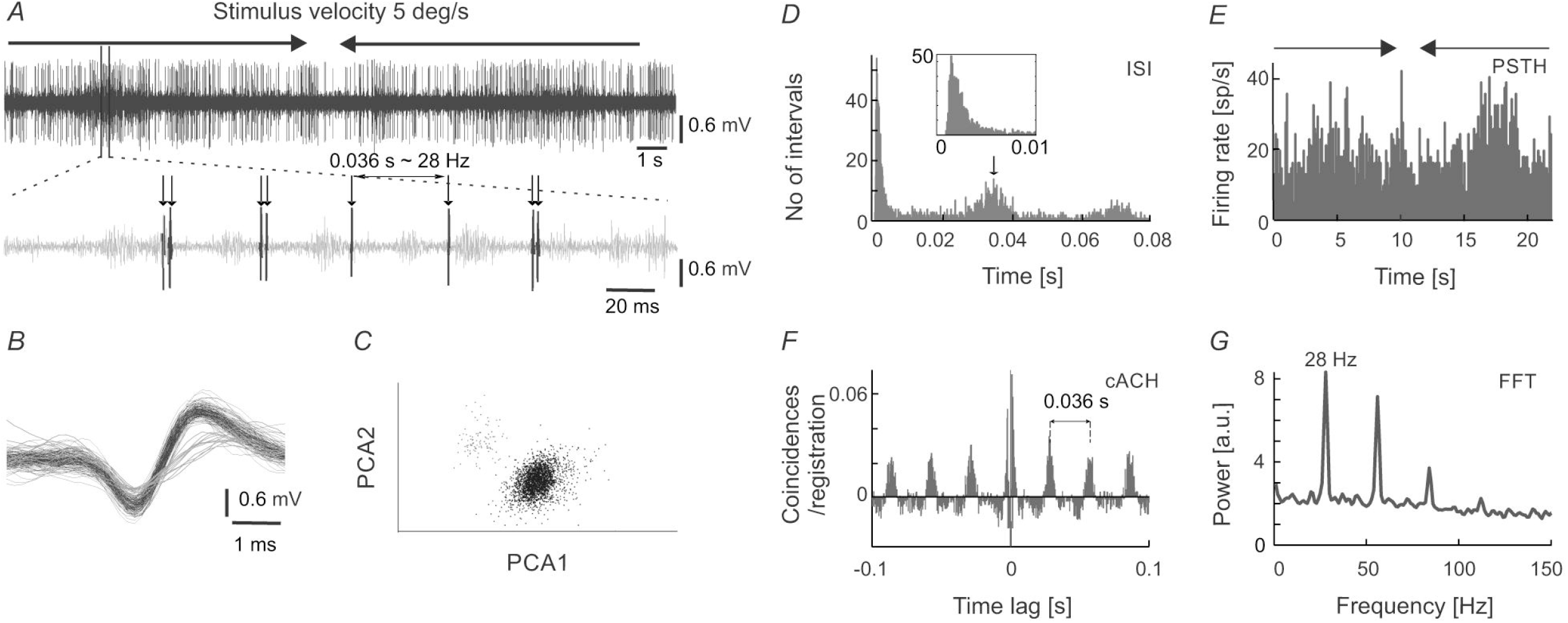
Example of strong similarity between frequencies of stimulus-phase-locked, stimulus-phase-independent, and background oscillations in SC neuron. (A), Raster plot showing stimulus-phase-locked oscillations. (B), PSTH of response in (A) (0.5 ms resolution), an average of 100 trials. (C), ACH from PSTH in (B) revealing stimulus-phase-locked oscillations at 10 Hz. (D), cACH after removing shift predictor showing the presence of oscillations at a similar frequency (cf. (C)). (E), Shift predictor of ACH in (C). (F), ACH of the background activity recorded after visual stimulation revealing a similar oscillatory frequency at 9.5 Hz. (G – J), Power spectra prepared for (C – F) respectively. Prominent peaks are visible at around 10 Hz in all power spectra.

A detailed analysis of the seven cells, for which all types of oscillations were observed, showed that in a majority of the recordings, the frequency of the stimulus-phase-locked oscillations corresponded to a subharmonic of the stimulus-phase-independent oscillatory frequency (e.g. Fig. 9). This kind of relationship is observed in different dominant frequencies for both types, but the frequency observed in the ACHPSTH and in the shift predictor (16.5 Hz; Fig. 10*G, J*) is half of the frequency observed in the cACH (33 Hz; Fig. 9*H*). It should be mentioned that the relationship between frequencies of different types of oscillations was not stable for a given neuron and could vary depending on stimulus parameters. In two (2 / 7) of recorded neurons, there was no clear relationship between the frequencies of stimulus-phase-locked (oscillations in the ACHPSTH and shift predictor) and stimulus-phase-independent oscillations (oscillations in the cACH; e.g. Fig. 10). Lack of relationship is revealed by the presence of different dominant frequencies observed in ACH_PSTH_ (Fig. 10*C, G*) and cACH (Fig. 10*D, H*).

**Figure 9.**
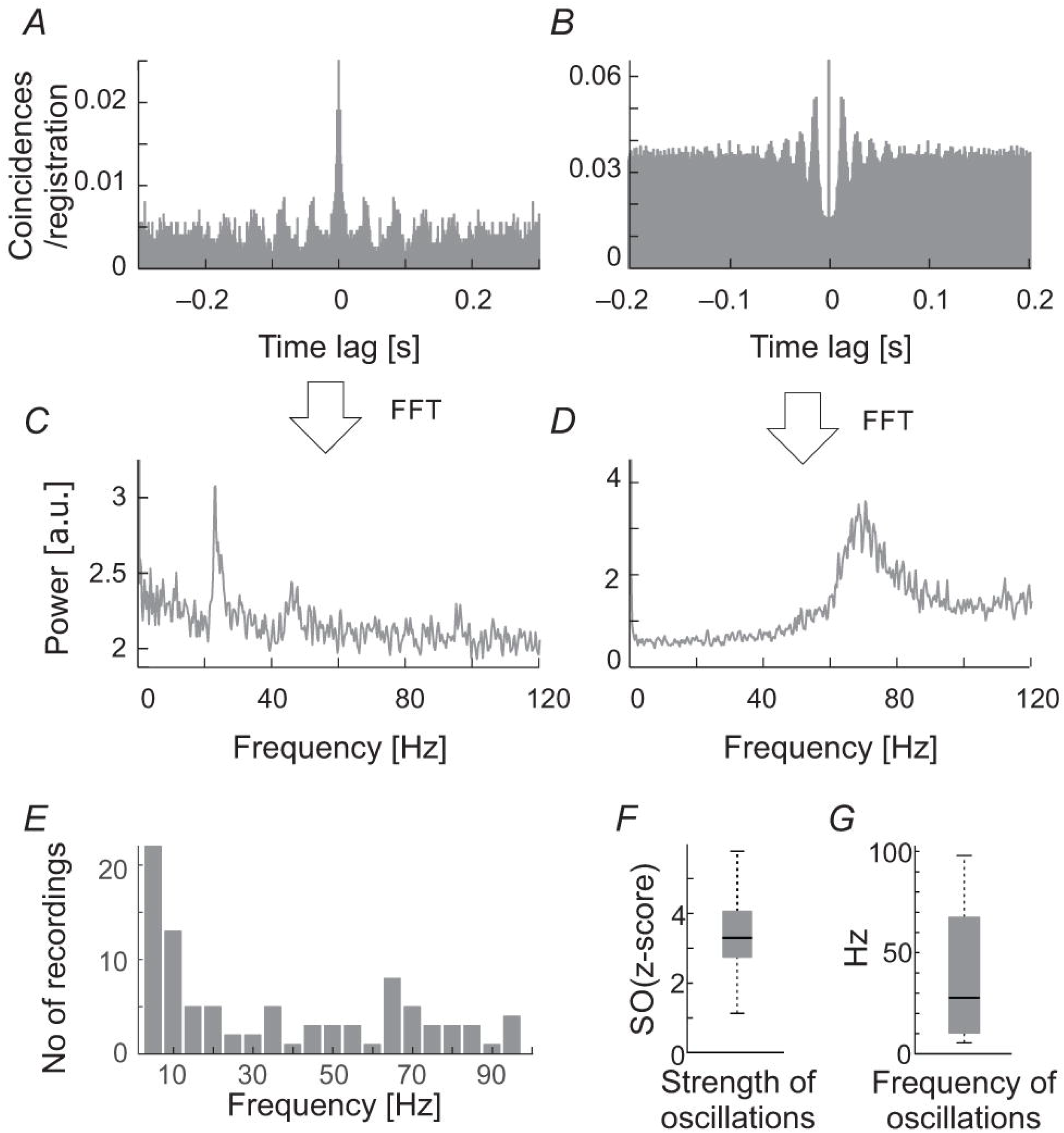
Relationship between stimulus-phase-locked, stimulus-phase-independent, and background oscillatory frequencies found in one neuron, showing stimulus-phase-locked oscillations as a subharmonic of oscillations. (A), Raster plot showing stimulus-phase-locked oscillations. (B), PSTH of the response shown in (A), averaged over 100 trials. The histogram was built with 0.5 ms resolution. (C), ACH prepared from PSTH (B), reviling stimulus-phase-locked oscillations in 16.5 Hz. (D), cACH after removing shift predictor, showing the presence of oscillations in similar frequency as in (C). (E), ACH prepared for background activity registered after visual stimulation, reviling similar frequency of oscillations as in previous diagram (33.3 Hz). (F), shift predictor for the same neuron indicating the same frequency as in (C). (G – J), Power spectra prepared for (C – F) respectively.

**Figure 10.**
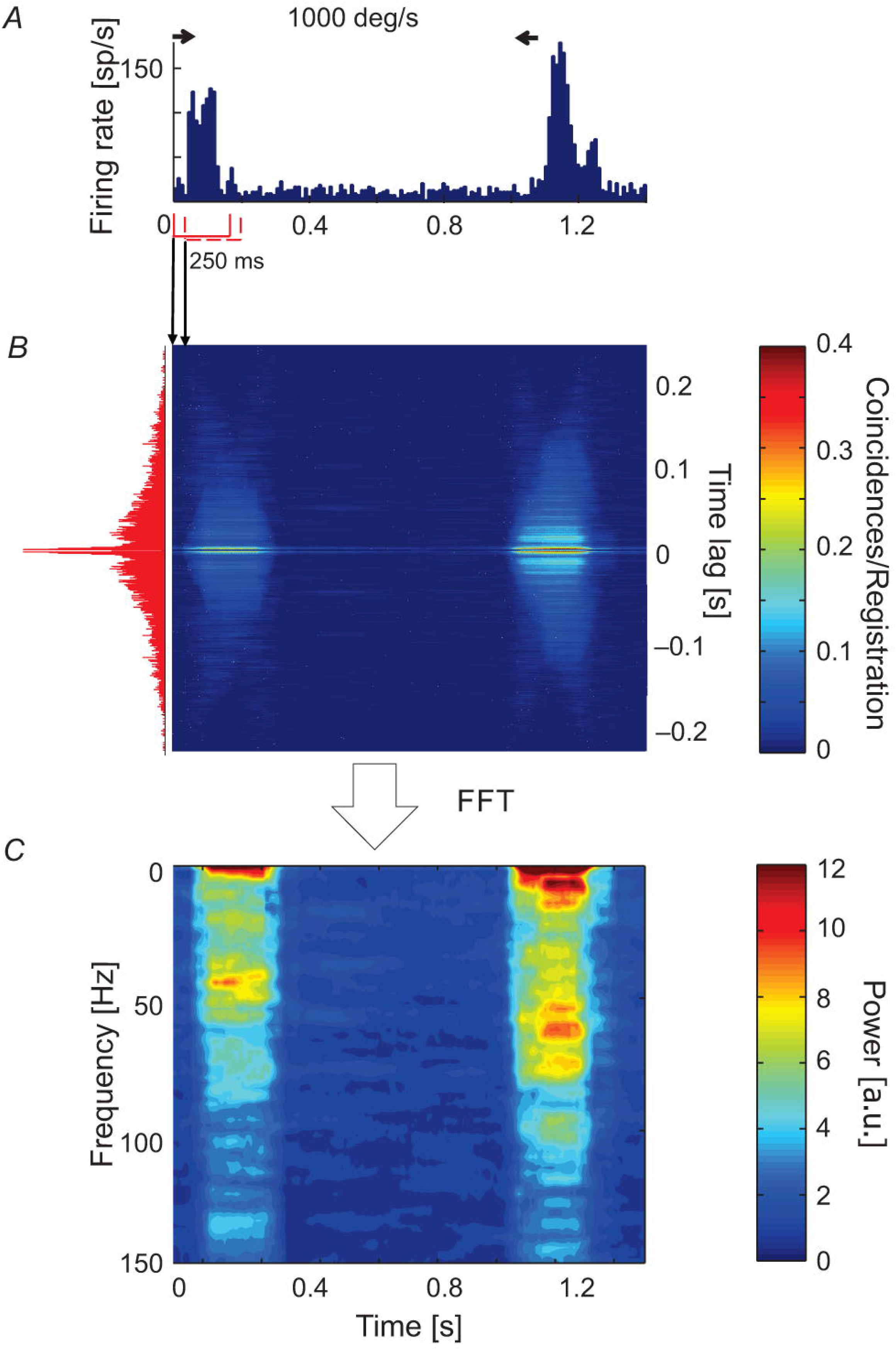
Relation of frequencies found as stimulus-phase-locked, stimulus-phase-independent and background oscillations for one neuron. (A), raster plot showing stimulus-phase-locked oscillations. (B), PSTH of response showed in (A), averaged over 100 trials (0.5 ms resolution). C, ACH_PSTH_ prepared from PSTH in (B), revealed stimulus-phase-locked oscillations at 10 Hz. (D), cACH after removing the shift predictor, showed the presence of oscillations at a much higher frequency (cf. (C)). (E), ACH of background activity after visual stimulation, revealed similar oscillatory frequencies as in panel (C). (F), Shift predictor for the same neuron indicating the same frequency as in (C). (G – J), Power spectra for (C – F) respectively.

### Renewal density analysis of neuronal activity in the SC

Renewal density analysis (RDA) was used in order to reveal the source of oscillatory activity (intrinsic or extrinsic - extracellular) (Mountcastle et al. 1990; Lebedev and Wise, 2000; Samonds and Bonds, 2005). In RDA method lower power spectrum value for dominant frequency may refer to the extrinsic source of oscillation. In case of no change in the power of oscillations in dominant frequency, we may assume that the oscillatory activity is a unique feature of recorded neuron. The temporal order of interspike intervals was destroyed by random shuffling and reconstructed surrogate spike trains were analyzed in the same way as the original. A comparison between surrogate (RDA) raw FFT analyses, both for spontaneous and visually evoked oscillations is shown in Figure 11 (Fig. 11*A* and *B*, and Fig. 11*D* and *F*, respectively, RDA FFT – black line and raw FFT - grey line). In the majority of cases, when strong oscillations in the beta/low gamma range were observed in background activity (e.g. Figure 11*A* and *B*), the peak of the dominant frequency decreased but did not disappear. In Figure 11*C* and *D* are shown examples of stimulus-phase-independent oscillations for which a large decrease in the amplitude of the dominant frequency is visible in the RDA FFT plots (Fig. 11*D*) indicating an extrinsic oscillatory source. Similar results were observed for stimulus-phase-locked oscillations (Fig. 11*E* and *F*).

**Figure 11.**
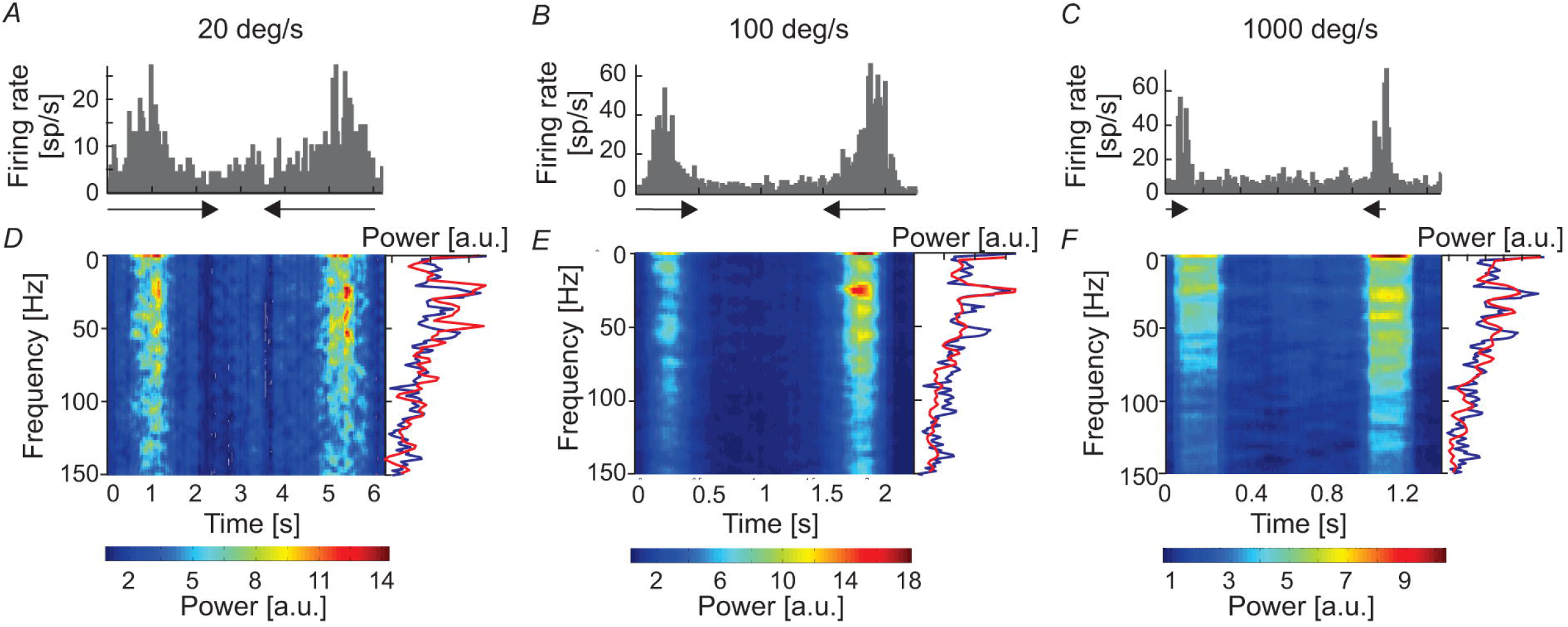
Determination of the origin of the three types of oscillatory activity using a renewal density analysis (RDA). (A) and (B), RAW FFT (grey lines) and RDA FFT (black lines) of neuronal background activity of two SC cells presented in Figure 2, a small decrease of peak amplitude in dominant frequencies in both cases (23 Hz and 70 Hz, respectively) does not suggest the extrinsic source of oscillations. (C) and (D), Example of ACH for raw data of neuron presenting oscillations classified as stimulus-phase-independent, and power spectra for raw data (grey line) and surrogate spike train containing shuffled inter-spike intervals (black line). The difference in power spectra of raw and surrogate data indicates the extrinsic source of oscillation in this neuron activity due to prominent peak disappearance (57 Hz). (E) and (F), Example of ACH for the neuron, which oscillatory activity was classified as phase-locked, and the related power spectra for the raw and surrogate spike trains. Complete peak disappearance in dominant frequency (20 Hz) in RDA FFT indicates an extrinsic source of the stimulus-phase-locked oscillations.

### Characteristics of stimulus-phase-locked and stimulus-phase-independent oscillations in the sSC

Figure 12 compares the characteristics of the two types of oscillation found in visually evoked activity for one direction of stimulus movement – to right. The results obtained for opposite stimulus direction were similar. The first row (Fig. 12*A, B*) shows the relationship between the frequency of the oscillations and stimulus velocity. A very weak and positive correlation was found between the frequency of the oscillations and stimulus velocity for stimulus-phase-independent and negative and much stronger correlation was found for stimulus-phase-locked oscillations. Lower frequencies were observed for slower vs. faster stimulus velocities (r = 0.15, p = 0.05; e.g. 40 (6) Hz at 5 deg s^−1^ vs. 56 (5) Hz at 1000 deg s^−1^; Mann-Whitney U-test, *p* = 0.041) in the case of stimulus-phase-independent oscillations. In contrast, the frequencies of stimulus-phase-locked oscillations in response to high stimulus velocities were significantly lower comparing to responses at low velocities (r = −0.36, p < 0.001; e.g. 38 (7.5) Hz at 5 deg s^−1^ vs. 20 (5) Hz at 1000 deg s^−1^; Mann-Whitney U-test, *p* = 0.022). There were no differences between frequencies of oscillations for the two movement directions for either type of oscillation (Wilcoxon test, phase-locked, *p* = 0.72; stimulus-phase-independent, *p* = 0.33). Panels *C - D* and *E - F* in Figure 12 present differences in the correlations between stimulus velocity and SO (quantified by SO(z-score) and Os) for the two types of oscillations. Independently of the measure, the strength of oscillations in the case of stimulus-phase-independent oscillations was not correlated with stimulus velocity for both movement directions (*r_R_* = 0.05, *p_R_* = 0.53; *r_L_* = 0.07, *p_L_* = 0.35 for Os index and *r_R_* = −0.08, *p_R_* = 0.29 and *r_L_* = −0.14, *p_L_* = 0.07 for SO(z-score) index, R and L indicate the direction of stimulus movement to the right or left, respectively). In contrast, both indices of oscillation strength in the case of stimulus-phase-locked oscillations were significantly correlated with stimulus velocity for both movement directions (*r_R_* = 0.61, *p_R_* < 0.001; *r_L_* = 0.59, *p_L_* < 0.001 for Os and *r_R_* = 0.44, *p_R_* < 0.001 and *r_L_* = 0.42, *p_L_* < 0.001 for SO(z-score)). Further analysis of the relationship between the SO(z-score) and the maximum firing rate in the response showed a weak correlation for stimulus-phase-independent oscillations (Fig. 12*G*; *r* = 0.16, *p* = 0.03), but a stronger correlation in the case of stimulus-phase-locked oscillations (Fig. 12*H*; *r* = 0.32, *p* < 0.001).

**Figure 12.**
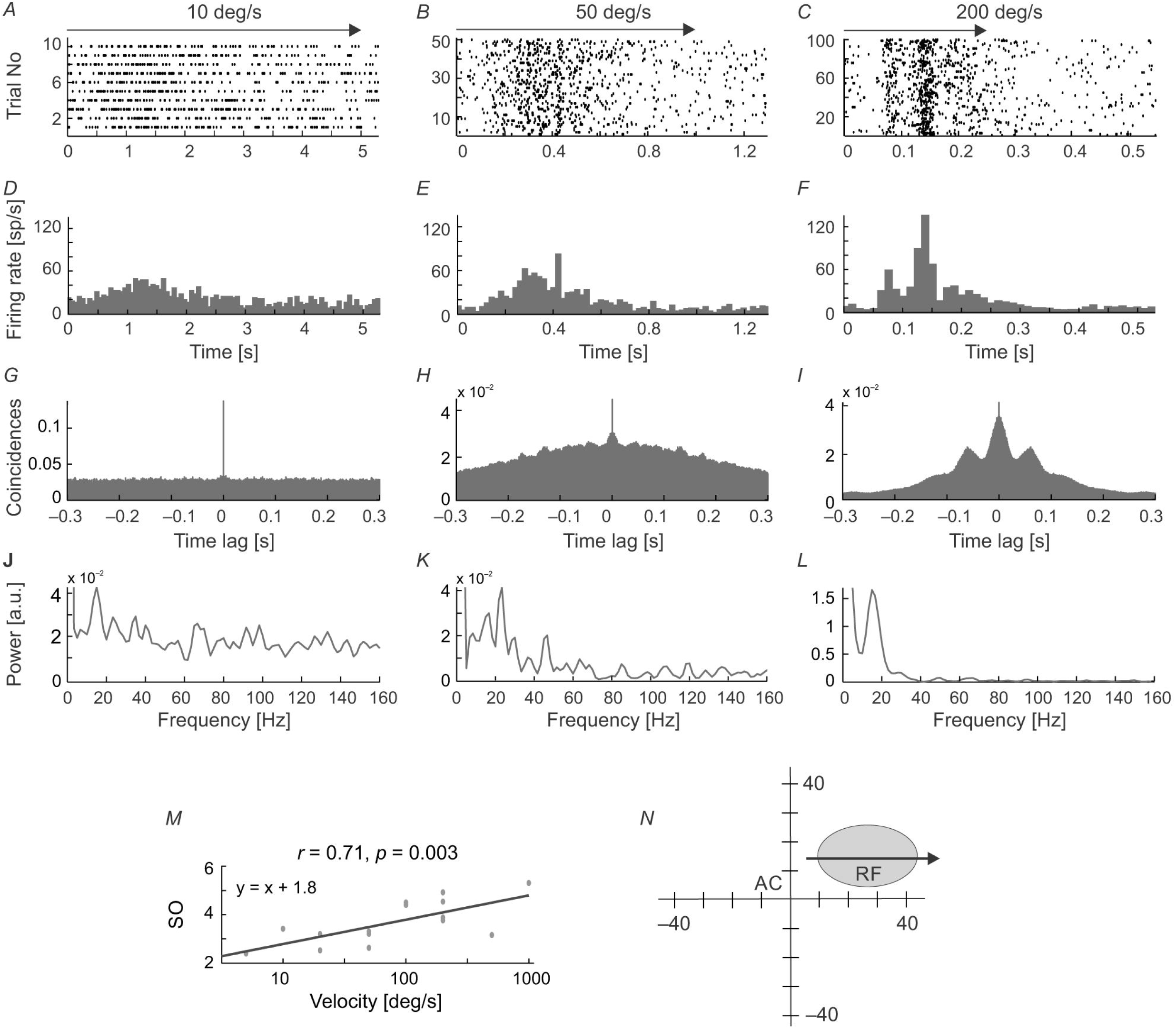
Summary results for all recorded SC cells and characteristics of stimulus-phase-independent oscillations (left column) and stimulus-phase-locked oscillations (right column) in evoked activity. (A), (B), The frequency range for the two types of oscillations for nine stimulus velocities tested. N - the number of neurons used in the calculations, n - the number of registrations used to do the calculations. (C – F), Correlations between the strengths of the oscillations and stimulus velocity measured by two coefficients, i.e. oscillation score (C, D) and SO(z-score) (E, F). (G), (H), Correlation between SO(z-score) and the maximum firing rate in the recorded responses. Maximum firing rate was measured as the maximum rate in the PSTH with the same number of bins (200) for each stimulus velocity. Note the stronger correlation for the phase-locked type of oscillation. (I), (J), Relationship between SO(z-score) and the frequency of the detected oscillations. (K), (L), Correlation between peak firing rate and frequency of oscillations. Note the large number of points below 40 Hz. Presented are results for one direction of stimulus movement – to right (away from *area centralis*). The results obtained for opposite stimulus direction were similar.

Stimulus-phase-independent oscillation frequencies (for all recordings; Fig. 12*I*) were distributed over the whole range of tested frequencies, while phase-locked oscillation frequencies (Fig. 12*J*) were mostly grouped below 40 Hz. For both types of oscillations, stronger oscillations were observed in lower frequencies (r = −0.26 for oscillations and r = −0.41 for stimulus-phase-locked oscillations, p < 0.001) The upper panel of Figure 13 shows the distribution of the main frequencies as histograms and raster plots (insets) in the background activity (Fig. 13A) and of stimulus-phase-independent and of stimulus-phase-locked oscillations detected in visually evoked activity (Fig. 13*B* and *C*). Stimulus-phase-locked oscillations occurred mostly in the low-frequency range (< 25 Hz), whereas the frequencies of stimulus-phase-independent oscillations, and those found in the background activity, were more uniformly distributed over the tested frequency range, albeit with some preference for low frequencies. We compared the characteristic frequencies of the different types of oscillations recorded from the same cell, in order to elucidate the relationship between oscillations found in visually evoked and background activity. Pairwise comparisons are presented in the lower panel of Fig. 13 (*D - F*). There was no difference between the distribution of characteristic frequencies of stimulus-phase-independent and background oscillations (cf. Fig. 13*A* and *B*; K-S test, *p* = 0.062; Mann-Whitney U-test, *p* = 0.54). The correlation analysis showed a strong linear dependence between characteristic frequencies of stimulus-phase-independent and background oscillations (Fig. 13*D*; *r* = 0.8, *p* < 0.001; e.g. in Fig. 4). However, there was a significant difference between distributions of frequencies of phase-locked and background oscillations (cf. Fig. 13*A* and *C*; K-S test, *p* = 0.001), and the correlation between detected frequencies of these oscillations failed to reach significance (Fig. 13*E*; *r* = 0.65, *p* = 0.055). Similarly, distributions of frequencies of stimulus-phase-locked and stimulus-phase-independent oscillations were significantly different (cf. Fig. 13*B* and *C*; K-S test, *p* < 0.001), and we have not found significant correlation between these frequencies (Fig. 13*F*; *r* = −0.1, *p* = 0.8; e.g. Fig. 8). These results indicate a strong similarity between stimulus-phase-independent and background oscillations and perhaps a common origin/mechanism. On the other hand, stimulus-phase-locked oscillations are closely related to fast changes in the receptive field and differ from the other types of oscillations in the SC.

**Figure 13.**
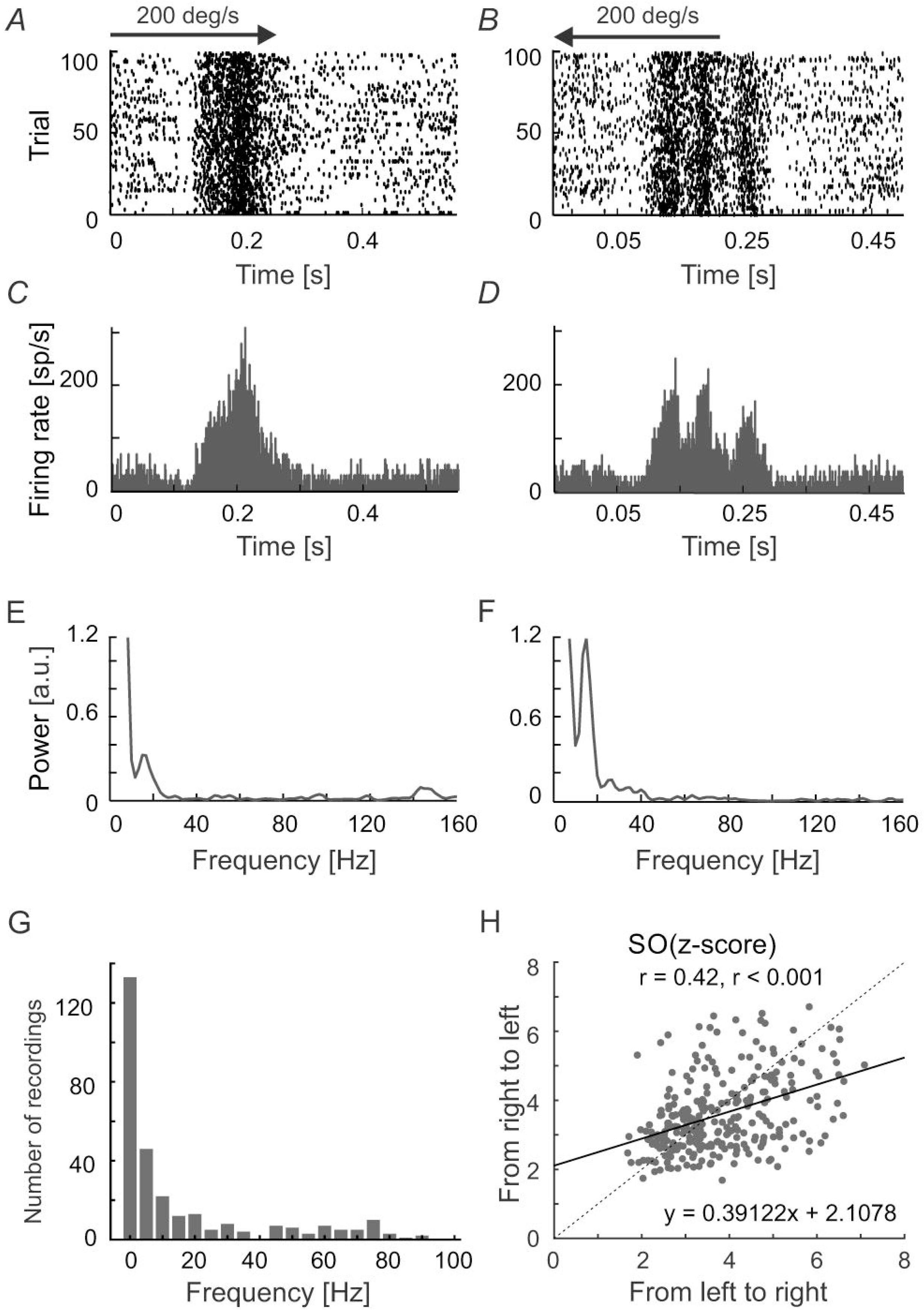
Comparison between the dominant oscillatory frequencies in the background, stimulus-phase-independent, and stimulus-phase-locked oscillations. (A - C), Distributions of the dominant frequencies for all types of oscillations found for individual neuronal activity. Inserts represent raster distributions from which histograms were prepared. (D), Pairwise comparison of frequencies found in sSC cell activity showing and background oscillations. (E), Comparison of frequencies found in single neuron activity showing both phase-locked and background oscillations. (F), Comparison of frequencies found in the activity of cells with stimulus-phase-locked and stimulus-phase-independent oscillations.

## Discussion

We confirm and broaden the previous reports on oscillations in spontaneous and visually evoked neuronal spiking activity in the superficial layers of the SC. Our studies indicate that stimulus-induced oscillations can appear in two forms – stimulus-phase-independent and stimulus-phase-locked to the stimulus onset. A comparison between the background and stimulus-phase-independent oscillation frequencies revealed that their frequencies were very similar, and suggests that stimulus-phase-independent oscillations may represent enhanced “background” oscillations. Stimulus-phase-locked oscillations were present in responses to flashing spots and to moving stimuli. Using quantitative measures of the strength of oscillations we showed a clear dependence between the strength of the stimulus-phase-locked oscillations and stimulus velocity – the higher the velocity, the higher the strength of the oscillations. We suggest that these oscillations are independent of oscillations present in spontaneous activity, although we cannot exclude the possibility that they result from a phase reset of the “background” oscillations.

### Oscillations in the background activity of SC cells

Our results indicate that oscillations in spontaneous activity are much more common than suggested by previous studies (Brecht et al. 1999; 2001; Chabli et al. 2000; Mandl, 1993). The origin of the spontaneous oscillations in SC has not been yet considered. We performed RDA (Mountcastle et al. 1990; Lebedev and Wise, 2000; Samonds and Bonds, 2005) to reveal the source (intrinsic - cellular or extrinsic - extracellular) of these “spontaneous” oscillations, which occur in a frequency range between 5 and 85 Hz. In earlier studies (e.g. Samonds and Bonds, 2005), any decrease in the amplitude of dominant frequency in surrogate spike train analysis was regarded as an indication of extrinsic oscillatory source. Our RDA results on the background activity may suggest an intra-collicular network or extra-collicular source for the oscillations due to the reduced peak at the oscillation frequency in the power spectrum (see below). However, a small reduction in the peak of the power spectrum could also occur if the oscillatory spiking activity is unstable during the recording epoch, which we think was the case in our recordings.

It has been suggested that the synchronous oscillatory activity in the LGN may be driven by retinal or cortical input (Ito et al., 2010). These two main sources of visual input to SC (for review see Waleszczyk et al. 2004; May, 2006) can be responsible, as well, for the presence of oscillations in spontaneous activity of SC cells. Simultaneous recordings from the retina and LGN in anesthetized cats studies by Neuenschwander and Singer (1996) reviled existence of stimulus-evoked correlated spiking activity in high gamma range typical for oscillations in retina and LGN (61-114 and 38-125 Hz, retina and LGN, respectively). Similarly, high oscillation frequencies were found in cat LGN cells recorded together with synchronized retinal input (S-potential in studies by Ito and co-workers; mean 79,9 Hz; Ito et al. 2010) and retino-thalamic synaptic potentials in whole-cell recording from relay thalamic neurons by Koepsell and coworkers (range 40-80 Hz; Koepsell et al., 2009). These studies strongly point for a retinal source of LGN oscillations in high gamma frequency band. What more, the LGN cells receive oscillatory input from retinal ganglion cells, even in the absence of visual stimulation (Koepsell et al., 2009) indicating that the origin of retinal oscillations is stimulus independent. Oscillatory activity in the LGN is present and even dominates, in the Y-type LGN cells (Ito et al., 2010). Thus, it is likely that Y-type retinal ganglion cells, many of which send axonal collaterals both to LGN and SC (Stone et al., 1979; Leventhal et al., 1985), may be the source of “spontaneous” oscillation in these two early visual structures. Indeed, in part of SC cells recorded in our study the frequency of spontaneous oscillations was in the high gamma range (Fig. 13A, e.g. Fig. 2 B). However, the majority of SC cells had oscillation frequencies in low gamma range, characteristic for cortical oscillations (Ito et al., 1994; 2010; Castello-Branco et al., 1998) and alfa-beta range (Fig. 13*A*, e.g. Fig. 2*A*). Clearly, further studies are needed to determine the origin of these background activity oscillations.

### Stimulus-phase-independent oscillations in the sSC

The origin of stimulus-phase-independent oscillations is most likely similar to that of the background activity due to the comparable frequencies of these oscillations in recorded collicular cells. Characteristic frequencies for these oscillations, which spanned the gamma band, were similar to those reported for the retina or LGN (Przybyszewski, 1993; Neuenschwander et al. 1996; Koepsell et al. 2009; 2010; Ito et al., 2010) and cortico–collicular and tecto-tectal synchronization (Brecht et al. 1998; 1999), and suggest an extrinsic (extracollicular) source. What more, similarly to observed in the present study collicular stimulus-phase-independent oscillations, retinal oscillations, and retinal origin LGN oscillations are not phase-locked to the stimulus onset (Neuenschwander and Singer, 1996; Koepsell et al., 2009), suggesting also the retinal origin of collicular stimulus-phase-independent oscillations. The results of the RDA for stimulus-phase-independent oscillations also suggest an extrinsic source for such oscillations.

The above mentioned earlier studies did not reveal any dependence between stimulus-phase-independent oscillations and such parameter of visual stimulation as stimulus movement velocity. Moreover, the dependence between the strength of the oscillation and firing rate has not been reported, although the dependence of the strength of oscillations on stimulus features (Engel et al., 1990; Gray et al., 1990) might suggest such dependence. In agreement with earlier studies, we also found that there was no relationship between the strength of stimulus-phase-independent oscillations and movement velocity or neuronal firing rate (see Figure 12).

Stimulus-induced phase-locking was recently shown in multi-unit and LFP activity of the ferret SC (Stitt et al. 2013). To our knowledge, there is no information concerning stimulus-induced phase-locking activity in single neurons of the cat’s SC. We show that stimulus-phase-locked oscillations in the SC is a phenomenon related to processing information about fast changes in the visual field (stimulus on-set and off-set or high stimulus velocity). This is manifested as a positive correlation between the strength of the oscillations and stimulus velocity, as well as a positive correlation between firing rate and oscillation strength (SO; Fig. 12). Changes in the response pattern due to different stimulus velocities were reported previously as ‘burst synchronization’ in the pre-trigeminal cat preparation (Mandl, 1993) and in anesthetized cats (Chabli et al. 2000). Our results also indicate that the oscillation-like temporal pattern of responses to spot movement and the strength of the stimulus-phase-locked oscillations can differ for the two directions of stimulus movement, which was not described, to our knowledge, in previous SC studies. What more, our results indicate that the strength of oscillations is greater for the direction in the visual field away from *area centralis*. Interestingly, the strength of neuronal responses in SC was reported to be also greater for the direction away from *area centralis* (Hoffmann, 1972; Schoppmann and Hoffmann, 1979), what agrees with a positive correlation between firing rate and oscillation strength found in our study. Different oscillations induced by different stimulation (for example different directions of stimulus movement) have been observed in the primary visual cortex (Friedman-Hill et al. 2000; Samonds and Bonds, 2005; Feng et al. 2010; Womelsdorf et al. 2012; Bharmauria et al., 2015; Bharmauria et al., 2016) and the premotor cortex (forearm movements; Lebedev and Wise, 2000). The majority of stimulus-phase-locked oscillations in our data were stronger for movement in the preferred direction (Fig 6). However, responses of a few cells were observed for which the strength of oscillations was lower for the preferred direction of stimulus movement. Interestingly, in this study, we observed stimulus-phase-locked oscillations in the case of SC neurons with the preference for the fast moving stimuli which are characterized by a low variability of visual responses from trial-to-trial (Mochol et al. 2010). It is very likely that stimulus-phase-locked oscillation is a factor which influence (increase) reliability of neuronal responses.

### Origin of stimulus-phase-locked oscillations in the sSC

Defining the origin of oscillatory SC activity is problematic due to the complex nature of the intra-SC anatomical network and connections with other brain areas. The suppression of firing rate at the beginning of some responses, in which strong stimulus-phase-locked oscillations are present (Fig. 5, 6), may suggest the involvement of intrinsic GABAergic neurons in the formation of this type of oscillation, such as a strong retinal input onto inhibitory SC interneurons (for rev. see Mize, 1992; Meredith and Ramoa, 1998; Schmidt et al. 2001; Kaneda et al. 2008). Lo and colleagues (1998) reported the presence of sodium spikes after a hyperpolarizing impulse in 96% of low-threshold spiking cells in SC slices (see also Edwards et al. 2002). The main feature of such neurons is that they can generate a rebound response in the form of bursts after depolarization to 47 – 54 mV (Lo et al. 1998). This bursting activity was characterized by frequencies similar to those observed in our data for stimulus-phase-locked oscillations and frequency of bursting activity was voltage dependent. This explanation would be consistent with the earlier description of stimulus-phase-locked oscillations as ‘burst synchronization’ (Mandl, 1993; Chabli et al. 2000).

Another possible source of phase-locked SC oscillations could include network connections with other structures or a phase-reset of ongoing activity. Connections between the mammalian SC and the parabigeminal nucleus (Sherk, 1979; Baizer et al. 1991; Harting et al. 1991b; Usunoff et al. 2007) is one such source. This modulation may be the effect of cholinergic inputs from the parabigeminal nucleus onto GABAergic interneurons, thus enhancing inhibition within the SC (Binns and Salt, 2000; Lee et al. 2001). Another possibility may be activity reverberations in colliculo-parabigemino-collicular loops. Oscillations in visually evoked LFPs in the rat and ferret SC have been observed in response to a flashing light (our unpublished results; Fortin et al. 1997; Stitt et al. 2013). Blocking synaptic transmission resulted in a decay of oscillatory activity and Fortin and colleagues suggested that stimulus-locked oscillations in the SC were an effect of processing information within the SC. Thus, it seems likely that the origin of stimulus-phase-locked oscillations is an effect of two phenomena, intrinsic neuronal properties and local network connections involving GABA-ergic neurons. Induced stimulus-phase-locked oscillations are not unique to the SC and were also found in the spiking (Gray et al. 1989; Gray and Singer, 1989; Bringuier et al. 1997; Friedman-Hill et al. 2000; Womelsdorf et al. 2012; Baranauskas et al. 2016; Le et al. 2017) or EEG activity (Tallon-Baundry et al. 1996) of the visual and auditory cortices (Noda et al. 2013).

It should be pointed out that presented here results were obtained in anesthetized animals and in fact type of oscillations, their frequencies, strength, and relations to visual stimuli may differ in the case of the healthy sleep and/or in the active state of the animal. It has been shown in the number of studies in sensory cortices of different modalities that brain state affects background and stimulus-evoked neuronal firing rate, temporal reliability of neuronal response, response latency and threshold of stimulus detection (Fazlali et al., 2016; Hasenstaub et al., 2007; Lee and Dan, 2012; Li et al., 2009; Haider et al., 2012; Niell and Stryker, 2010; Schröder and Carandini, 2014; Zhao et al., 2014). As concerns influence of the brain state on the response pattern to sensory stimulation, brain state modulates in a different way early and late phase of cortical neurons’ responses (Fazlali et al., 2016). Although this effect was not studied in details, Fazlali and colleagues have shown the appearance of the stimulus-phase-locked oscillatory pattern in the late phase of the response of neuron in the somatosensory cortex in the desynchronized brain state (Fazlali et al. 2016, Fig. 3). Further studies are needed to elucidate whether similar phenomena is present in the subcortical structures of the visual system, particularly in the sSC.

The question arises as to whether the SC oscillatory activity is merely an epiphenomenon or reflects the oscillatory activity of its input structures (retina, visual cortex, Neuenschwander and Singer, 1996; Neuenschwander et al., 1999; Koepsell et al., 2009), and/or has a specific function in visual perception. One possibility is “binding coding” or synchronization (for reviews, Singer, 2001; Klimesh et al. 2007) in which oscillations are a basic mechanism of network cooperation and the transfer of information to other areas. We hypothesize that stimulus-dependent, and in particular, stimulus-phase-locked oscillations can enhance information transfer by increasing the probability of spike generation in the SC neurons’ projection targets. Stimulus-phase-locked oscillations most-likely also increase the reliability of neuronal responses in the SC. Further studies are needed to confirm these hypotheses.

## Additional Information

### Competing interest

**None.**

### Author contribution

W.J.W. and A.G designed the experiments; A.T.F., A.G., and W.J.W. performed the experiments; A.T.F. analyzed the data; A.T.F., A.G., and W.J.W. wrote the manuscript.

### Funding

The work was supported by grant from Polish Ministry of Science and Higher Education and National Science Centre N N303 820640.

## Acknowledgments

We would like to thank Marek Wypych and Gabriela Mochol for participation in some experiments and Joanna Smyda for technical assistance.

Current address of A. Foik: Department of Anatomy and Neurobiology, School of Medicine, University of California. Medicine Surge II, Room 364, Irvine 92697, USA. Current address of A. Ghazaryan: Institute of Applied Problems of Physics NAS RA, 25 Hr. Nersisyan St., 0014 Yerevan, Armenia.

## Abbreviations

ACH: autocorrelation histogram
cACH: corrected autocorrelation histogram
DiI: 1,1’-dioctadecyl-3.3,3’,3’-tetramethylindocarbocyanine perchlorate
ECoG: electrocorticogram
FFT: Fast Fourier Transform
ISI: interspike interval histogram
LGN: lateral geniculate nucleus
PSTH: peri-stimulus time histogram
r: Pearson correlation coefficient
rACH: raw autocorrelation histogram
RDA: renewal density analysis
RF: receptive field
SC: superior colliculus
sSC: superficial layers of superior colliculus
std(FFT): standard deviation of power spectrum

